# Comparative genome microsynteny illuminates the fast evolution of nuclear mitochondrial segments (NUMTs) in mammals

**DOI:** 10.1101/2023.04.20.537758

**Authors:** Marek Uvizl, Sebastien J. Puechmaille, Sarahjane Power, Martin Pippel, Samuel Carthy, Wilfried Haerty, Eugene W. Myers, Emma C. Teeling, Zixia Huang

## Abstract

The escape of DNA from mitochondria into the nuclear genome (nuclear mitochondrial DNA, NUMT) is an ongoing process. Although pervasively observed in eukaryotic genomes, their evolutionary trajectories in a mammal-wide context are poorly understood. The main challenge lies in the orthology assignment of NUMTs across species due to their fast evolution and chromosomal rearrangements over the past ∼200 million years. To address this issue, we systematically investigated the characteristics of NUMT insertions in 45 mammalian genomes, and established a novel, synteny-based method to accurately predict orthologous NUMTs and ascertain their evolution across mammals. With a series of comparative analyses across taxa, we revealed that NUMTs may originate from non-random regions in mtDNA, tend to locate in transposon-rich and intergenic regions, and unlikely code for functional proteins. Using our synteny-based approach, we leveraged 630 pairwise comparisons of genome-wide microsynteny and predicted the NUMT orthology relationships across 36 mammals. With the phylogenetic patterns of NUMT presence-and-absence across taxa, we constructed the ancestral state of NUMTs given the mammal tree using a coalescent method. We found support on the ancestral node of Fereuungulata within Laurasiatheria, whose subordinal relationships are still controversial. This strongly indicates that NUMT gain-and-loss over evolutionary time provides great insights into mammal evolution. However, we also demonstrated that one should be cautious when using ancestral NUMT trees to infer phylogenetic relationships. This study broadens our knowledge on NUMT insertion and evolution in mammalian genomes and highlights the merit of NUMTs as alternative genetic markers in phylogenetic inference.

## Introduction

Nearly all eukaryotic nuclear genomes have constantly been challenged by the insertion of foreign DNA of various origins, essentially shaping their architecture over evolutionary time (Adams et al. 2000; Richly and Leister 2004; Kleine et al. 2009). In mammalian genomes, the acquisition of extrinsic DNA is largely conducted by the horizontal transfer of mitochondrial DNA (mtDNA) segments, leading to nuclear pseudogenes of mitochondria origin (NUMTs) (Mourier et al. 2001; Timmis et al. 2004; Hazkani-Covo et al. 2010). Although the mechanism of this ongoing process is not yet fully understood, the well-accepted hypothesis proposes that the passive uptake of mtDNA fragments into the nuclear genome occurs at double-stranded DNA breaks (DSBs) via non-homologous end-joining (NHEJ) repair machinery (Blanchard and Schmidt 1996). A few lines of evidence have suggested the association between NUMT integrations and human diseases (Turner et al. 2003; Wei et al. 2022); however, they are commonly regarded as ‘dead on arrival’ pseudogenes, as evidenced by the presence of stop codons, indels and frameshifts caused by random mutation, and the differences in the genetic code between the nuclear genome and mitogenome (Perna and Kocher 1996). While arguably not functional, NUMTs have been responsible for many occasions of misinterpretations in mtDNA heteroplasmy detection (Albayrak et al. 2016), mitochondrial disease studies (Wallace et al. 1997; Yao et al. 2008) and phylogenetic placements (Sorenson and Quinn 1998; Thalmann et al. 2004; Li et al. 2016), due to their homology to mtDNA. Attempts should, therefore, be made to identify NUMTs in genomes in order to avoid erroneous conclusions in mtDNA-related research.

NUMTs have been studied in a wide range of mammals, but discrepancies in their radiation, genomic distribution, mtDNA origin, functionality, and insertion time and rates were reported (Hazkani-Covo et al. 2010). The genetic basis underlying these variations and NUMT evolutionary trajectories in a mammal-wide context are poorly understood. It has been recently suggested that NUMTs, the molecular fossils of ancestral mtDNA, can be potential genetic markers to infer phylogenetic relationships (Bensasson et al. 2001). However, the application is only limited to a few studies that focused on groups of species in narrow phylogenetic brackets, such as Primates (Hazkani-Covo and Graur 2007; Hazkani-Covo 2009), Passeriformes (Liang et al. 2018) and Chiroptera (Puechmaille et al. 2011). The major challenge resides in the assignment of NUMT orthology across mammals, owing to the rapid gain and loss of NUMTs (Hazkani-Covo et al. 2010), fast sequence changes (Hazkani-Covo et al. 2010), and considerable chromosome rearrangements over 200 million years of evolution (Pevzner and Tesler 2003). Currently, the primary method to predict orthologous NUMT loci across species is by means of aligning NUMT sequences along with their flanking regions or, through whole genome alignments (Hazkani-Covo et al. 2010). These alignment-based approaches are only feasible to predict NUMT orthology within closely-related species in which the non-coding genomic regions are well aligned (Cunningham et al. 2022). Hence, to elucidate NUMT evolution in mammals it is imperative to develop new methods that enable NUMT orthology assignment between distantly-related species.

Genomic synteny has been deeply conserved across the tree of vertebrates (Simakov et al. 2020). Microsynteny is defined as a fine-scale genomic region in which the order of a number of genes is evolutionarily conserved across species (Kawashima 2019). It provides a valuable framework to interpret gene orthology relationships between species, especially for large multigene families or fast-evolving non-coding genes such as NUMTs (Kawashima 2019). In this study we comprehensively investigated the characteristics of NUMT insertions in 45 mammalian nuclear genomes and established a novel, synteny-based approach to accurately predict orthologous NUMTs and ascertain their evolution across mammalian clades. We observed that the Primate suborder Haplorhini has undergone a burst of NUMT insertions, while multiple NUMT expansion events may have occurred during the evolution of marsupials. Using comparative analyses across taxa, we showed that the mtDNA regions from which NUMTs originate are non-random. We also showed that NUMTs are likely to locate in transposon-rich and intergenic regions, and unlikely code for functional proteins. Using the novel approach we established, we performed 630 pairwise comparisons of genome-wide microsynteny and assigned NUMT orthology relationships across 36 mammals. We further constructed the ancestral state of NUMTs using a coalescent approach and discovered that the phylogenetic patterns of NUMT presence-and-absence in Laurasiatheria support the ancestral clade of Fereuungulata. This challenges previous phylogenies which placed bats as the sister clade to ungulates, but agrees with the recent genome-based topologies which support a sister-group relationship between carnivores and ungulates. These results indicate that NUMT gain-and-loss over evolutionary time can provide great insights into mammal evolution. However, we also demonstrated that one should be cautious when using ancestral NUMT trees to infer phylogenetic relationships. This study deepens our understanding of NUMT insertion and evolution in mammalian nuclear genomes and highlights the merit of NUMTs as alternative molecular markers in phylogenetic inference.

## Results

### Overview of NUMT insertions in mammalian genomes

Using the optimal methods (see Methods), we obtained a landscape of NUMT insertions across 45 mammalian genomes (Supplemental Tables S1-4; Supplemental Fig. S1). The number of NUMT insertions ranges from 43 (manatee; *Trichechus manatus*) to 3,886 (Tasmanian devil; *Sarcophilus harrisii*), with the median of 218 (Fig. 1a). Our predictions are comparable to the numbers reported in previous studies (Hazkani-Covo et al. 2010; Hazkani-Covo 2022). Despite large variation in numbers across species, NUMTs only represent, on average, less than 0.01% of a genome (Fig. 1a). The genome of the Tasmanian devil (*S. harrisii*) carries the longest cumulative NUMT length (1,951 kb) while the shortest was found in the common shrew *Sorex araneus* (15.07 kb). The individual NUMT length varies between 30 bp and 16,699 bp across species, and exhibits a similar distribution in species within the same order (Kolmogorov-Smirnov tests; Supplemental Figs. S2a, S2b). 20 out of 45 genomes were found to possess exceptionally long NUMTs (over 10 kb), some of which were derived from almost the entire mtDNA (Supplemental Table S2). Interestingly, the species in Cetartiodactyla have the highest percentages of complex NUMT blocks which comprise multiple individual NUMTs located within a genomic distance of 2 kb (see Methods; Supplemental Table S2). To demonstrate the reliability of predicted NUMT blocks, as an example we validated the genomic locus of the largest NUMT block in the *Molossus molossus* genome using our published PacBio raw reads for the genome assembly (Supplemental Fig. S2c). We noticed that NUMT content illustrates a strong positive correlation with genome size and genome transposable element (TE) content (specifically LINEs) respectively (*P* < 0.05; Spearman’s correlation tests), but significances of the tests dropped after the phylogeny correction (*P-adj* > 0.05; Fig. 1b; Supplemental Fig. S3). In addition to the similar NUMT length distribution observed among closely-related species, these results suggest that NUMT insertions, to some degree, follow an evolutionary pattern and could be potential genetic markers for phylogenetic inference.

**Figure 1:**
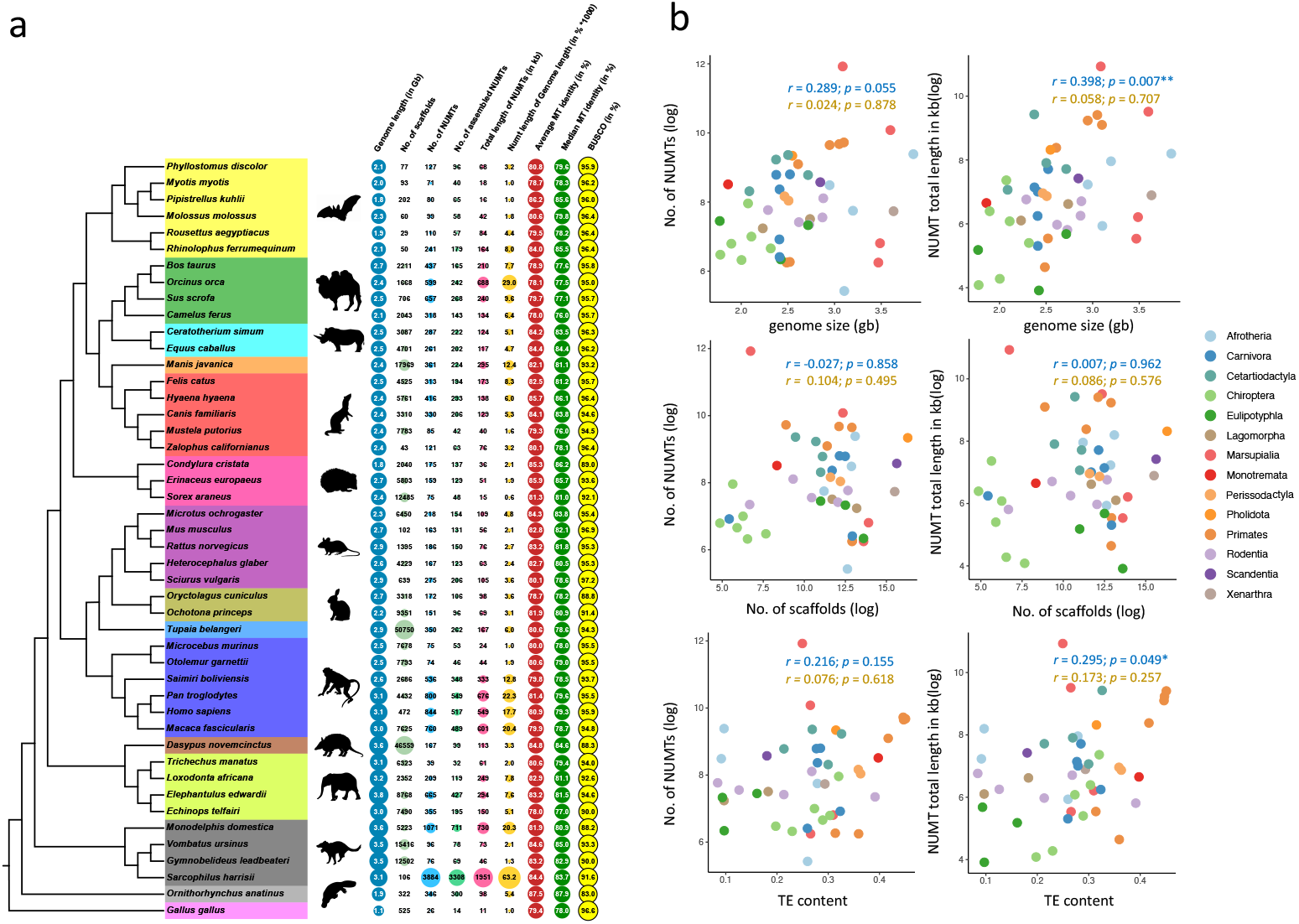
Overview of NUMTs in 45 mammalian genomes. **a)** Genome statistics and NUMT statistics in 45 mammalian species and the outgroup *Gallus gallus*. The phylogenetic relatedness of these 46 species was obtained from (Jebb et al. 2020) and the species highlighted in the same colour are in the same order. The columns, from left to right, represent genome length (Gb), number of scaffolds, number of NUMTs, number of assembled NUMT blocks, cumulative NUMT length (kb), fraction (% × 1000) of total NUMT length relative to genome length, average and median NUMT sequence identity to the corresponding mtDNA respectively, and BUSCO scores (%). **b)** Correlation between genome statistics and NUMT statistics. The scatterplots show the relationship between genome statistics (genome size, number of scaffolds, and average transposable element (TE) content) and NUMT statistics (number of NUMTs and accumulative NUMT length; log_2_ transformed). Correlation coefficients (*r*) and *P*-values were computed using Spearman’s correlation tests. In the scatterplots, coefficients (*r*) and *P*-values in blue and gold indicate the values before and after phylogeny correction (*0.01 < *P* < 0.05) (see Methods). The colour code indicates the species from the same order.

### NUMT expansions in primate and marsupial species

We observed a burst of NUMT insertions in Primates and Marsupialia. In Primates, four species with large NUMT numbers (547 ∼ 846) lead to the node of the suborder Haplorhini, whereas the other two species studied in the sister suborder, Strepsirrhini, only have a small number of NUMT insertions (76 ∼ 77) (Fig. 1a). This suggests that a burst of NUMT insertions occurred in Haplorhini, after its divergence with Strepsirrhini. In Marsupialia, similar to our observation on the Tasmanian devil (*S. harrisii*), a recent study found that two species in the family Dasyuridae, the Tasmanian devil (*S. harrisii*) and the yellow-footed antechinus (*Antechinus flavipes*), had rapid NUMT expansions (Hazkani-Covo 2022). However, we also revealed that the opossum (*Monodelphis domestica*) in the family Didelphidae underwent a similar burst of NUMTs (1,083) as the Tasmanian devil, contrary to the possum *Gymnobelideus leadbeateri* (76) and the common wombat *Vombatus ursinus* (112) in the family Petauridae and Vombatidae, respectively (Fig. 1a). This result indicates that massive expansions of NUMT content may have occurred multiple times during the evolution of marsupials, and a thorough taxonomic sampling is crucial to drawing accurate conclusions on NUMT expansions.

### The mtDNA regions from which NUMTs originate are non-random

It was reported that certain regions in mtDNA, such as the D-loop, tend not to produce NUMTs in a few primate genomes (Tsuji et al. 2012), while it is well-represented by NUMTs in the cow and a few cetacean genomes (Ko et al. 2015; Grau et al. 2020). By scanning the mitogenome with a 50 bp sliding window for each species (320 windows representing 16,000 bp in mtDNA per species), we conducted comparisons of the coverage between all possible windows across 45 species using pairwise Mann-Whitney *U* tests (Figs. 2a-c; see Methods). We observed that the distribution of the NUMT coverage varies across windows (Fig. 2a). Of all 51,040 comparisons, 9,273 (18.17%) tests yielded significant results (FDR < 0.01; Fig. 2b), and some windows (e.g. w1∼50; w200∼277) illustrate different distributions of coverage from others (Fig. 2c). To further verify these results, we simulated coverage data for the null distribution of equal coverage by randomly reshuffling mtDNA coordinates of NUMTs for each species, performed pairwise comparisons across windows, and repeated these analyses 1,000 times. We found that the number of significant tests yielded from the observed data (9,273) is approximately 10 times as many as that of the simulated data (median: 953, Supplemental Fig. S4a). This result supports our hypothesis that the mtDNA regions from which NUMTs originate are non-random. In addition, at single-nucleotide resolution the mtDNA coverage of NUMTs exhibits a species-specific pattern across taxa (Supplemental Fig. S4b).

**Figure 2:**
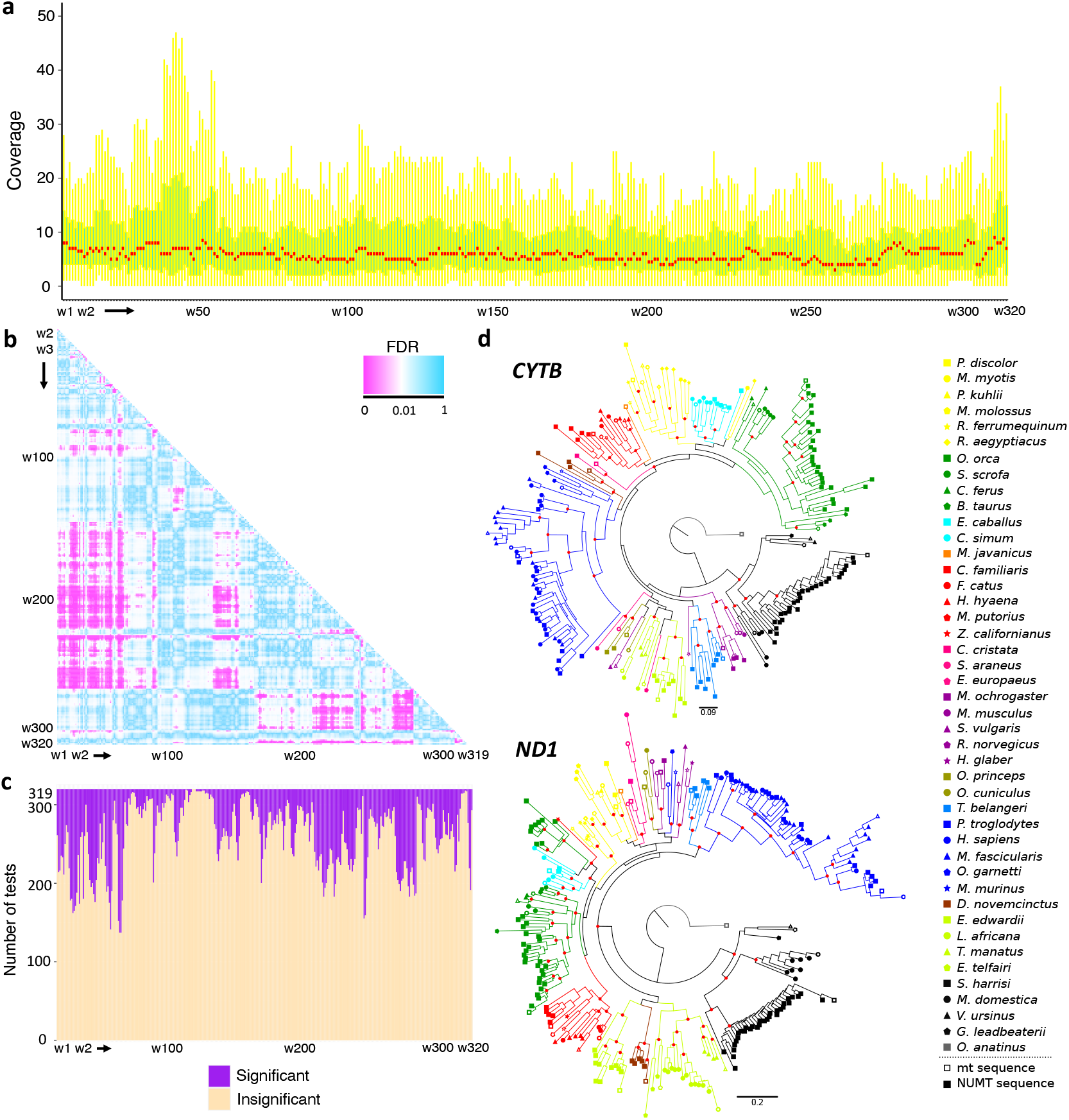
The mtDNA origin of NUMT sequences and mammalian NUMT phylogeny. **a)** Distribution of mtDNA coverage by NUMTs in a 50 bp window resolution. The *x*-axis represents 320 windows and the *y*-axis represents the coverage distribution of each window across 45 species. The median value of each window is highlighted in red while the interquartile range is in green. The outliers are not shown in the boxplot. **b)** Pairwise comparisons of mtDNA coverage by NUMTs between 50 bp sliding windows using Mann-Whitney *U* tests. 320 windows (w1 ∼ w320) representing 16,000 bp in mtDNA were investigated. The heatmap depicts the FDR values of 51,040 tests between all possible windows. Pink colours indicate that the tests are significant (FDR < 0.01), while blue colours indicate that the tests are insignificant (FDR ≥ 0.01). **c)** Distributions of significant and insignificant testing results from pairwise comparisons between all possible windows. The coverage of each window across species was compared to those of the rest 319 windows respectively. **d)** Mammalian NUMT phylogeny. Two maximum-likelihood phylogenetic trees were inferred using 188 NUMT sequences mapped to the *Cytb* locus and 215 NUMT sequences mapped to the *ND1* locus across species, respectively. For each species, the mtDNA *Cytb* and *ND1* loci were also included to infer the two NUMT trees, respectively. The effective alignment lengths for the phylogenetic inference are 1,472 bp (*Cytb* locus) and 1,174 bp (*ND1* locus), respectively. The red dots on the nodes of the trees indicate high branch supports (HS-aLRT > 90%, aBayes > 0.9 and UFBoot > 90%). Mitochondrial sequences in each species are indicated by hollow symbols.

### NUMT phylogenetic trees largely agree with the mammal phylogeny

To investigate the evolution of NUMTs, we constructed two maximum-likelihood phylogenetic trees using the NUMT sequences that mapped onto mtDNA *CYTB* and *ND1* loci respectively, across 45 mammals (see Methods). Both NUMT trees, to a broad extent, agree with the mammal phylogeny (Jebb et al. 2020), in which Monotremata, Marsupialia and Eutheria clades are supported (Fig. 2d). For the *CYTB* NUMT tree, the major disagreements lie in the fact that odd-toed ungulates were positioned within bats, armadillo was a sister group to primates, and lagomorphs together with squirrel were placed within Afrotheria (Fig. 2d). In contrast, for the *ND1* NUMT tree armadillo was placed within Afrotheria, bats were split into two clusters which are the sister groups to Ungulata and Eulipotyphla respectively, and odd-toed ungulates were positioned within even-toed ungulates (Fig. 2d). These results suggest that NUMTs are under limited selective pressure during their evolution.

### NUMTs tend to locate in transposon-rich and intergenic regions

Some previous evidence suggests that NUMTs appear to insert into the genomic locations with rich transposable element (TE) content (mainly retrotransposons) (Tsuji et al. 2012). To further examine this association across mammals, we firstly investigated the TE content in 5kb flanking regions (both upstream and downstream) of each NUMT/NUMT block (see Methods) with a window size of 500 bp. Although exhibiting different distributions across orders (Supplemental Fig. S5a), the TE content in the 5 kb flanking regions of NUMTs is significantly higher than the TE genome content in 30 out of 45 mammals (*P* < 0.05; Mann Whitney *U* tests), except some species belonging to rodents and marsupials (Supplemental Fig. S5b). This indicates that a burst of NUMTs in *S. harrisii* and *M. domestica* was driven by rapid insertion rather than duplications of pre-existing NUMTs. When analysing windows across species, we observed that all the windows had a significantly higher TE content than the genome average TE content (*P* < 0.05), except the first upstream and downstream windows (Mann-Whitney *U* tests; Fig. 3a). These two windows, directly connecting with NUMTs, had a significantly lower TE content than the remaining flanking windows up to 5 kb at both ends across species (*P* < 0.05, Mann-Whitney *U* tests; Fig. 3b). Aligned with the previous study (Tsuji et al. 2012), we postulated that NUMTs tend not to directly insert into TEs. This hypothesis is further supported by our observation that newly inserted NUMTs are located closer to TE than older NUMTs, as evidenced by a negative correlation between NUMT/mtDNA sequence identity and the distance of NUMTs to their closest TE in over two thirds of mammals (see Methods; Spearman’s correlation tests; Supplemental Fig. S6). All these results indicate that NUMTs have a tendency to locate in genomic regions adjacent to TE, which may lead to the subsequent expansion of NUMT content in genomes.

**Figure 3:**
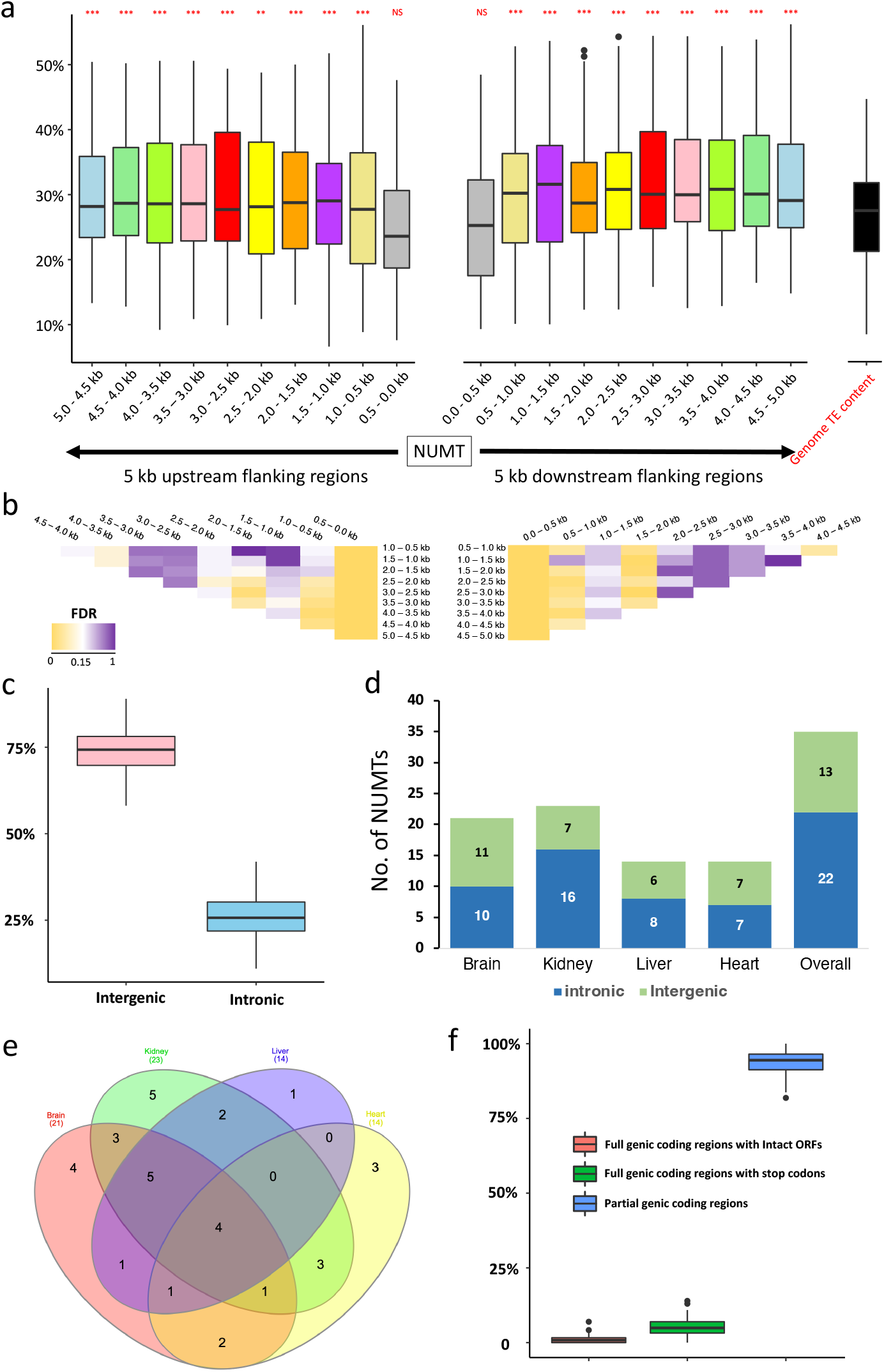
Characteristics of mammalian NUMTs. **a)** Transposable element (TE) content in the 5kb up- and down-stream of flanking regions of NUMTs/NUMT blocks with a window size of 500 bp. For each species, the median TE content was estimated for each of 20 windows in the flanking regions of all the NUMTs/NUMT blocks. The black boxplot on the right indicates the average genome TE content in 45 mammals. The TE content of each window was respectively compared to the average genome TE content using Mann Whitney *U* tests (upper-tailed, paired mode; *0.01 < *P* < 0.05; **0.001 < *P* < 0.01, \*\*\**P* < 0.001). **b)** Pairwise comparisons of TE content across all windows in the flanking regions of NUMTs/NUMT blocks. The heatmap illustrates the significance (corrected by FDR) of the Mann-Whitney *U* tests (yellow: low; white: median; purple: high). The comparisons were performed separately for the windows from the 5kb up- and down-stream of flanking regions. **c)** Distribution of genomic loci (intergenic and intronic) in which NUMTs were identified across 36 mammalian genomes. **d)** Distribution of intergenic and intronic NUMTs/NUMT blocks that are expressed in brain, liver, kidney and heart samples from platypus. **e)** Venn diagram showing the number of expressed NUMTs shared amongst brain, liver, kidney, and heart samples from platypus. **f)** The boxplots indicate the distribution of individual NUMTs that contain intact ORFs, full genic coding regions with stop codons, and incomplete genic coding regions across 45 species.

Next, we investigated if NUMTs are prone to locate in introns or intergenic regions. Across the 36 mammals investigated, the number of NUMTs detected in intergenic regions was significantly higher than intronic regions, with the ratio approximately 3 to 1 (*P* < 0.001, Mann-Whitney *U* test; Fig. 3c). The pale spear-nosed bat (*Phyllostomus discolor*) has the highest percentage of intergenic NUMTs (89%), whereas the highest percentage of intronic NUMTs (41.9%) is observed in the platypus (*Ornithorhynchus anatinus*). While it was estimated that the cumulative sizes of introns and intergenic regions in animal genomes are similar (Francis and Worheide 2017), our result indicates that NUMTs tend to locate in intergenic regions.

### Most NUMTs are not expressed and non-functional

Upon integration into nuclear genomes, NUMTs are generally considered non-functional (Leister 2005). To confirm this hypothesis, we employed stringent criteria (e.g. junction expression between a NUMT and its flanking regions) to examine NUMT expression in four tissue types across five species (Supplemental Fig. S7a; see Methods). We found that NUMTs are rarely expressed, with a small percentage (0.15% ∼ 4.51%), on average, expressed across different tissues of these species (Supplemental Fig. S7b). Using the platypus as an example, we observed that expressed NUMTs are significantly enriched in introns (*P* = 0.0136, Chi-square test; Fig. 3d). We speculated that intronic NUMTs can be co-expressed with their host genes without *de novo* innovation of independent promotors. This hypothesis may also explain the high percentage of expressed NUMTs seen in platypus (Supplemental Fig. S7b) as its genome has the highest percentage of intronic NUMTs. It is noteworthy that, using our method we also identified polymorphic NUMTs that are individual specific in platypus (Supplemental Fig. S7c), which is commonly seen in mammalian genomes (Dayama et al. 2014; Dayama et al. 2020; Wei et al. 2022). In addition, we noticed that 4 NUMTs were expressed across all four tissue types in platypus, suggesting that they are under some selective constraints (Fig. 3e). However, we found that on average 99.2% of NUMTs across species contain incomplete open reading frames (ORFs) of mtDNA genes or full genic coding regions with stop codons, based on the nuclear genetic code (Fig. 3f; Supplemental Table S5). Despite being expressed, they are unlikely to be translated into proteins or, at least, not functioning as protein-coding genes.

### NUMT presence-and-absence patterns are alternative molecular markers to infer mammal phylogeny

To investigate the evolutionary trajectories of NUMTs across mammals, we developed a novel method that utilises protein-coding genes in a conserved genomic synteny block as anchors to locate NUMTs and analyses their mtDNA origin to infer NUMT orthology between each pair of species (see Methods; Supplemental Fig. S8a). The estimation of error probability and expectations (*E*) indicates high accuracy and reliability of our method (see Methods; Supplemental Figs. S8b, S8c). By leveraging 630 pairwise comparisons of genome microsynteny among 36 mammals, we observed that only a small proportion of NUMTs/NUMT blocks were predicted to be orthologous in each species, except human (*Homo sapiens*), chimpanzee (*Pan troglodytes*) and macaque (*Macaca fascicularis*) (Fig. 4). A relatively large number of orthologous NUMTs/NUMT blocks were found in primates, carnivores, bats, and ungulates, while only a few were identified within the remaining defined clades (see Methods; Supplemental Fig. S9). Using phylogenetic patterns of NUMT presence-and-absence, we employed a coalescent approach to predict the NUMTs that are ancestral to each node given the mammal tree (see Methods). The number of ancestral NUMTs/NUMT blocks predicted on the node decreases as the divergence time increases (Fig. 5a), suggesting that NUMT gain and loss follows an evolutionary pattern and could infer mammal phylogeny. Unsurprisingly, 258 ancestral NUMTs/NUMT blocks were predicted on the node branching to human, chimpanzee, and macaque that diverged only 29 Mya (Jebb et al. 2020). 23, 22, 18 and 11 NUMTs/NUMT blocks were, respectively, found ancestral to Carnivora, Cetartiodactyla, Chiroptera, and Primates (Fig. 5a). As the similar divergence time as the above orders in Laurasiatheria, the root of Rodentia was predicted to possess no ancestral NUMTs (Fig. 5a). This is possibly due to the fact that rodent species have undergone a high level of genome reshuffling, whose rates are much greater than other mammalian orders such as Carnivora and Primates (Capilla et al. 2016). These arrangements, such as DNA insertions, inversions and translocations, disrupt the analogy of genome organisation across species so that our microsynteny-based method is not powerful enough to identify orthologous NUMTs that are located in highly reshuffled regions. Markedly, 7 orthologous NUMTs/NUMT blocks were identified in the species across the defined clades (see Methods; Fig. 5a, Supplemental Table S6). Two NUMTs (Candidates 1 and 2) were regarded ancestral to Boreoeutheria, which were predicted in 17 species across 5 orders and 15 species across 6 orders, respectively (Fig. 5a). We also identified one NUMT (Candidate 3) shared by 17 species that lead to the ancestor of Eutheria, but did not find any NUMTs ancestral to the root of mammals (Fig. 5a). These results are not surprising because, as relics of ancient mtDNA, NUMTs evolve under limited selective constraints (Bensasson et al. 2001). Excitingly, we noticed that the phylogenetic patterns of Candidates 4, 6 and 7 support the ancestral clade of Fereuungulata within Laurasiatheria (Fig. 5a), strongly indicating that NUMT presence-and-absence can provide great insights into mammal phylogeny.

**Figure 4:**
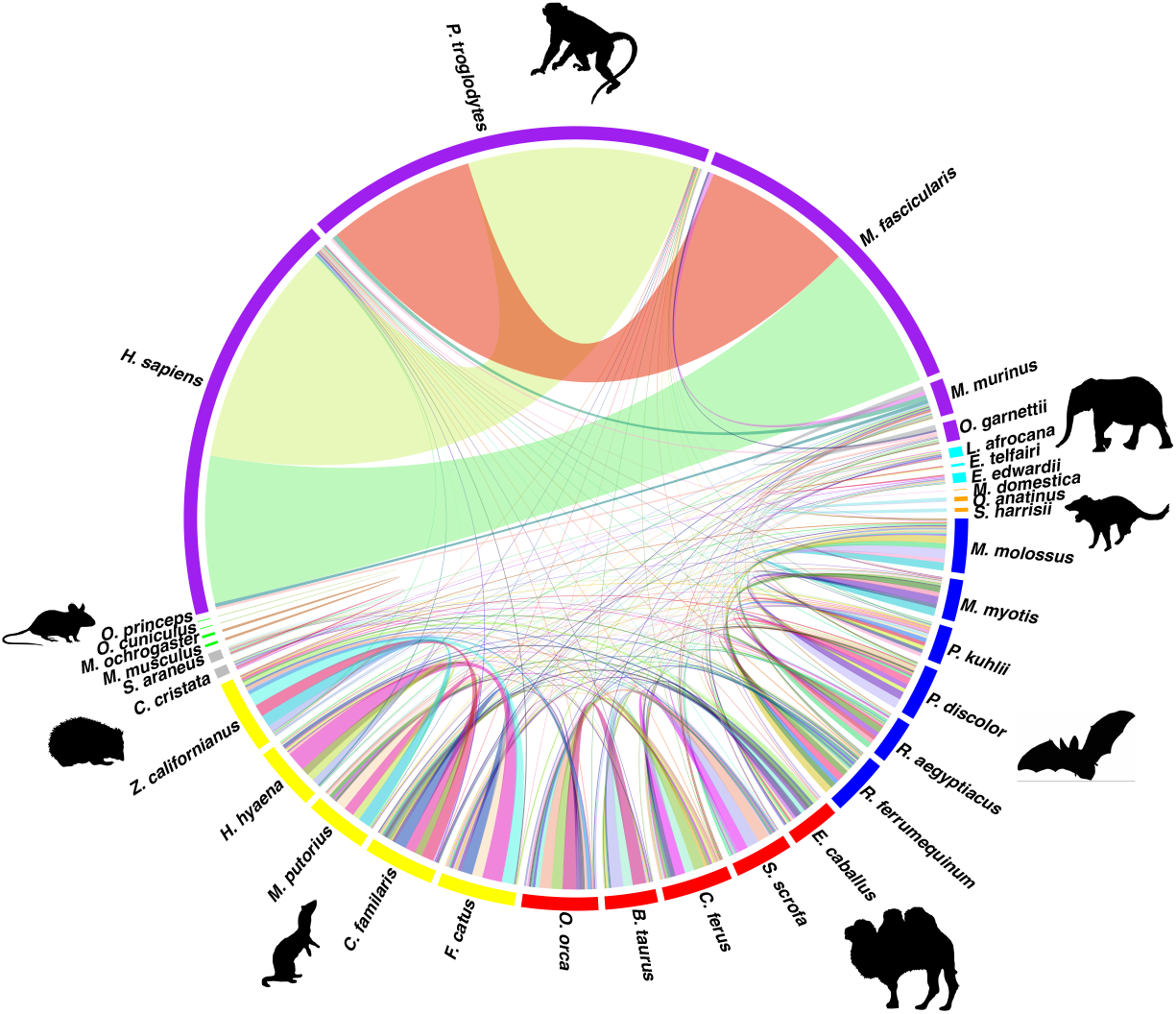
Circos diagram showing the number of predicted orthologous NUMTs/NUMT blocks among 36 species in a pairwise manner. The width of the link between two species indicates the number of orthologous NUMTs/NUMT blocks predicted. The colour code of the outside layer indicates the species in different defined clades (see Methods).

**Figure 5:**
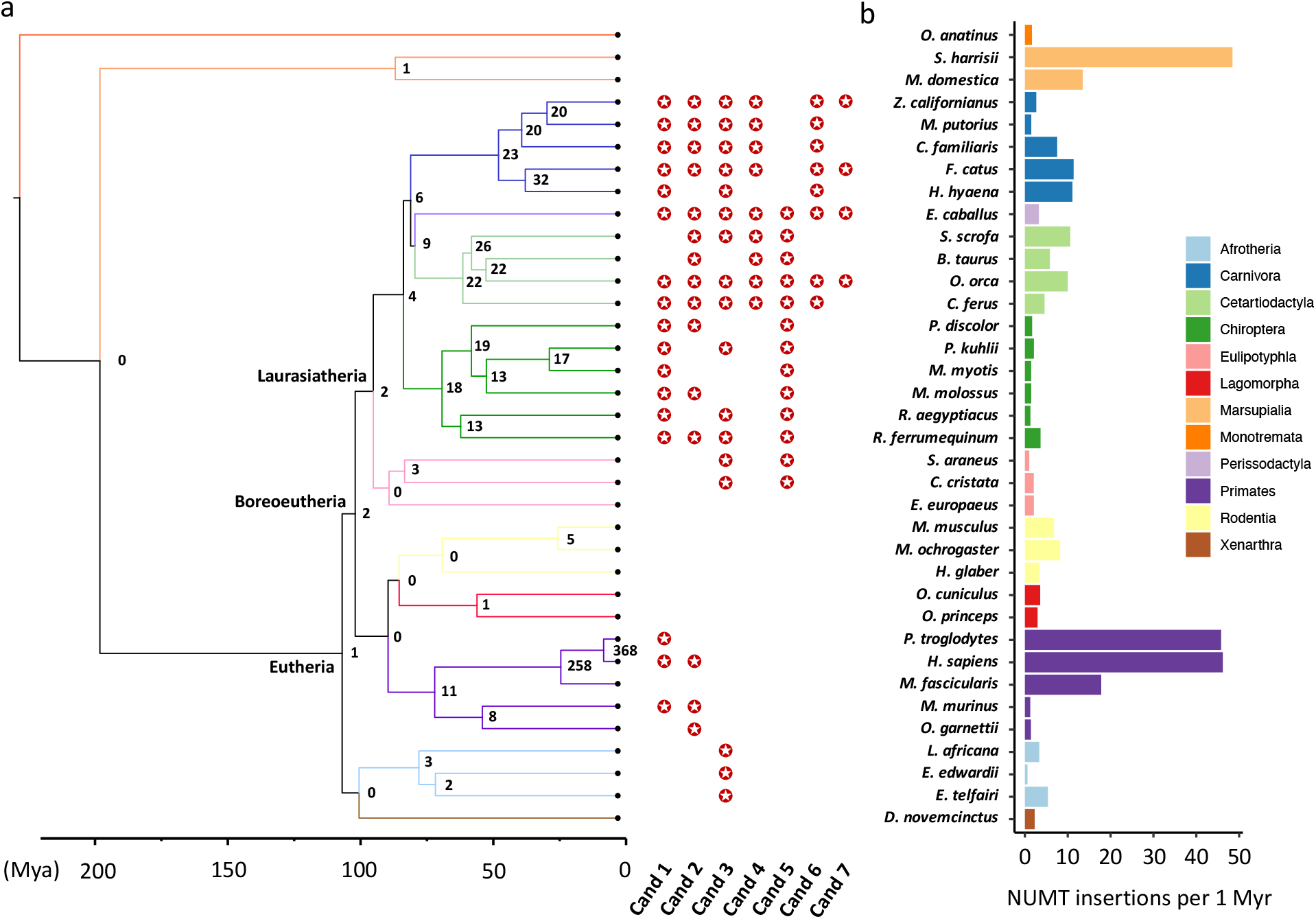
Ancestral NUMTs across mammalian clades and NUMT insertion rate. **a)** The numbers of ancestral NUMTs/NUMT blocks on the nodes of the phylogenetic tree. The numbers were inferred on the basis of the predicted NUMT orthology relationships across 36 mammals using a coalescent approach. Seven NUMTs/NUMT blocks (Cand 1∼7) that were predicted as orthologs in the species across the defined clades are highlighted. **b)** The estimation of NUMT insertion rates (number of NUMT insertions per 1 Myr) across mammals. For each species, the rate was estimated by dividing the number of individual, non-orthologous NUMTs, the individual NUMTs not found in its closely-related species or monophyletic group, by the divergence time (see Methods). The colour-code in both panels indicates the species in the orders they belong to.

### Human, chimpanzee, and Tasmanian devil genomes exhibit the highest NUMT insertion rates

Little attention has been paid to explore NUMT insertion rates in mammals due to the difficulty of NUMT orthology assignment across species. Using the NUMT orthology relationships we predicted and the divergence time within mammals, we estimated the NUMT insertion rate for each species (see Methods; Supplemental Table S7). We observed that the species within Marsupialia (Tasmanian devil) and Primates (human and chimpanzee) have higher insertion rates compared to the remaining species (Fig. 5b). An earlier study predicted that NUMT insertion rates in human and chimpanzee were 5.7 and 7.7 NUMTs per 1 Myr respectively (Hazkani-Covo and Graur 2007), whereas we estimated much higher insertion rates (46.2 and 45.8 NUMTs per 1 Myr for human and chimpanzee respectively; Fig. 5b). The disparities lie mainly in the fact that we predicted nearly twice as many NUMTs (846 in human and 819 in chimpanzee; Fig. 1a) as the numbers in that study (452 and 469 in human and chimpanzee respectively (Hazkani-Covo and Graur 2007)), and a higher percentage of species-specific NUMTs (27.4% as opposed to 15% (Hazkani-Covo and Graur 2007)) whose insertions are deemed to occur after their speciation.

### Cautions should be taken when using ancestral NUMT trees to infer phylogeny

To see if alignments of ancestral NUMTs are appropriate to infer phylogenetic relationships, we used a maximum-likelihood method to construct phylogenetic trees for three orthologous NUMTs (Candidates 1-3) predicted across distant orders (see Methods). We observed that the trees inferred from Candidates 1 and 2 are highly similar to the mammal tree (Jebb et al. 2020), with most species in the same order grouped together (Fig. 6). The exceptions lie in *Microcebus murinus* and a few bat species in Candidate 1 and *Equus caballus* in Candidate 2. For the tree inferred from Candidate 3, the two main clades Boreoeutheria and Atlantogenata are unambiguously split but the phylogenetic relatedness within Boreoeutheria is largely unresolved (Fig. 6). Although NUMT evolution largely agrees with our understanding of mammal phylogeny (Fig. 2d), phylogenies inferred from ancestral NUMTs should be interpreted with caution.

**Figure 6.**
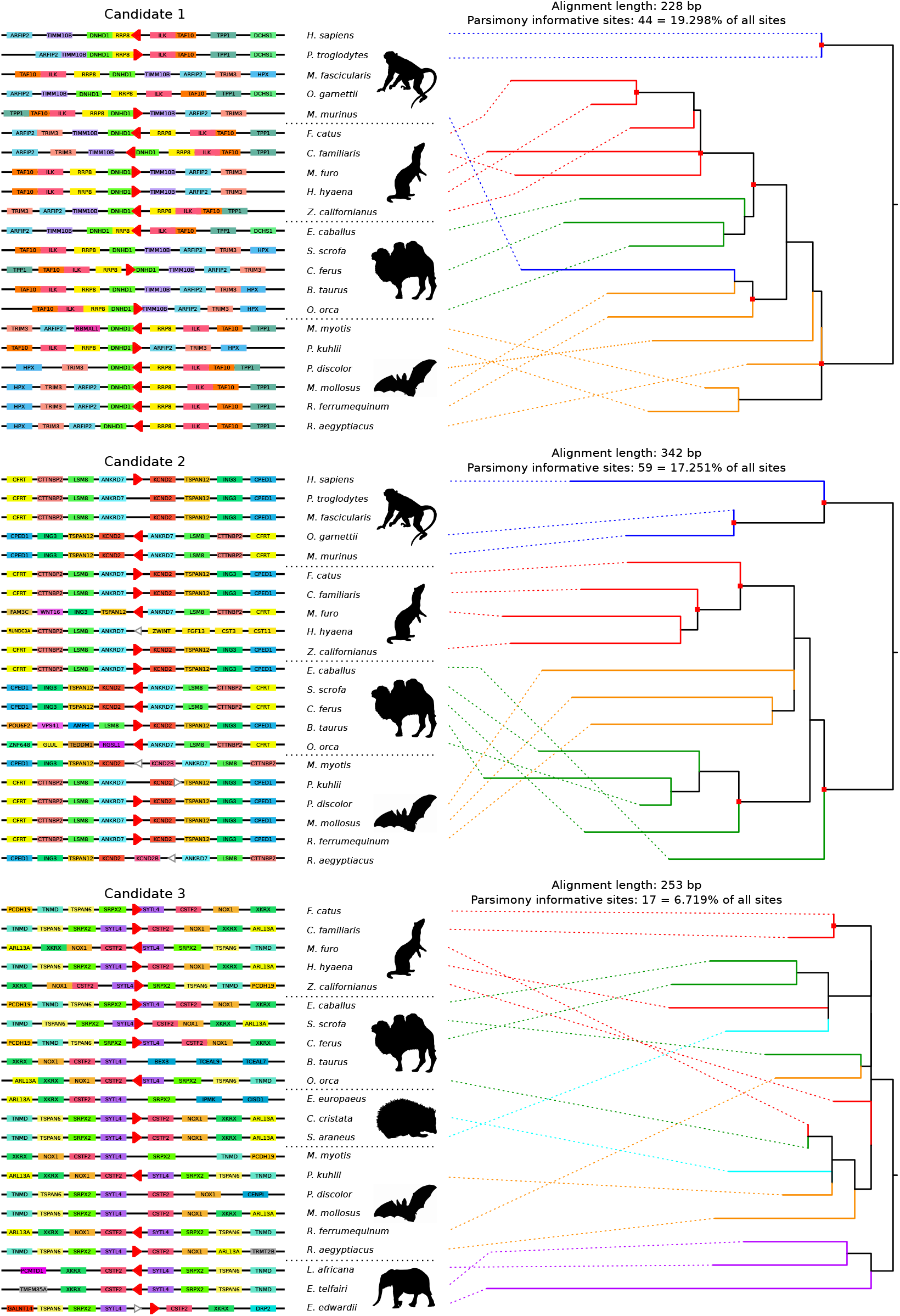
Visualisation of the genome microsynteny of 3 ancestral NUMTs (Candidates 1-3) predicted across the defined clades and their phylogenetic trees. For each candidate, the lines represent the conserved microsynteny blocks across species. Protein-coding genes are shown as rectangles on the lines, which are colour-coded. Genes with no gaps between each other indicate that they have overlapping genomic coordinates. NUMTs are represented by triangles, with its direction relative to the corresponding mtDNA indicated. Red triangles represent ancestral orthologous NUMTs, while white ones represent NUMTs that are considered as non-orthologous. The triangles that overlap with protein-coding genes indicate that the NUMTs are located in the intronic regions of these protein-coding genes; otherwise, the NUMTs are located in intergenic regions or 3’-UTRs. The phylogenetic tree for each NUMT candidate was constructed using a maximum likelihood approach. The effective alignment length and the number of parsimony informative sites for each tree are indicated on the plot. The red dots on the nodes of the trees indicate high branch supports (HS-aLRT > 90%, aBayes > 0.9 and UFBoot > 90%). The colour code of tree branches indicates the species in the defined clades, and the dashed lines connect the species and the branches they belong to.

## Discussion

In this study, we systematically investigated the characteristics of NUMT insertions in 45 mammalian genomes, and determined their radiation, genomic distribution, mtDNA origin, functionality, and insertion rates across species. Using a novel, synteny-based approach we established, we leveraged 630 pairwise comparisons of genome-wide microsynteny to ascertain NUMT evolution across mammals. We demonstrated that some lineages in Primates and Marsupialia exhibit a burst of NUMT insertions (Fig. 1a). However, the underlying reasons for these expansions are currently not well understood. It is speculated that the increase of NUMT insertion can be attributed to changes in environmental factors (Wang et al. 2012). It has been proven that yeast cultured under non-optimal temperatures demonstrated an accelerated rate of mtDNA escape to the nucleus genome (Cheng and Ivessa 2012). For marsupials, it was hypothesised that the expansion of NUMTs in Dasyuridae may result from the rapid drop of global temperature, an event known as the Miocene climate transition that occurred shortly after the divergence of Dasyuridae (Kealy and Beck 2017). However, this conjecture is weakened by the paucity of NUMTs found in species with a similar divergence time.

By analysing NUMT coverage in mtDNA with a 50 bp sliding window, we revealed that the mtDNA regions from which NUMTs originate are not random (Figs. 2a-c). It will be interesting to see if the current conclusion stands when more species are included in the analyses. Additionally, at single-nucleotide resolution the coverage of NUMTs in mtDNA exhibits a species-specific pattern. Consistent with the previous study (Tsuji et al. 2012), we noticed that some regions within the D-loop, mainly the heavy strand region, seldom produce NUMTs in most of the species in Primates (Supplemental Fig. S4b). Nevertheless, a large percentage of NUMTs derived from mtDNA D-loop was observed in several species, such as cat, rat, American pika, and Tasmanian devil, representing different phylogenetic orders (Supplemental Fig. S4b). Our results suggest that the NUMT origination from mitogenomes is not completely stochastic, and also support the current hypothesis that NUMT insertion results from the degradation of abnormal mitochondria in which mitogenomes are randomly sheared (Kleine et al. 2009). It is noteworthy that we analysed the NUMT coverage in mtDNA up to 16,000 bp, which only covers part of the D-loop region. Due to the complex features of the D-loop (e.g. length differences across species; species-specific tandem repeats), the NUMT coverage in the D-loop region needs to be further explored.

We also demonstrated that NUMTs tend to locate in transposon-rich and intergenic regions (Figs. 3a-c). Unlike transposable elements, NUMTs do not have the mechanisms to duplicate and translocate themselves independently, and ample studies (Tsuji et al. 2012; Michalovova et al. 2013; Wang et al. 2020), including our analyses (Supplemental Fig. S3), reported a strong positive correlation between NUMT content and genome TE content. This suggests that NUMT expansions are associated with transposon activities. In addition, it is important to note that the genomic regions into which NUMTs insert might be random, but these regions may be immediately subject to certain selective constraints, such as purifying selection, so that newly-inserted NUMTs may be quickly eliminated. Therefore, our result is not surprising because introns are broadly more functionally conserved than intergenic regions due to the enrichment of *cis*-regulatory elements in introns, such as intronic splicing enhancers (ISEs) and silencers (ISSs) (Chorev and Carmel 2012; Shaul 2017). Disruptions of these regulators and exon-intron splice junctions via NUMT insertion may alter gene expression or produce dysfunctional proteins, leading to detrimental consequences (Turner et al. 2003; Goldin et al. 2004).

It is methodologically challenging to assign NUMT orthology across mammals owing to the fast evolution of NUMTs and their flanking regions, so that traditional alignment-based approaches are not feasible (Hazkani-Covo 2009; Hazkani-Covo et al. 2010). To address this challenge, we established a novel, synteny-based and accurate method to assign NUMT orthology relationships between species. By constructing the ancestral state of NUMTs, we revealed that the presence-and-absence patterns of 3 NUMT blocks (Candidates 4, 6, 7) support the ancestral clade of Fereuungulata (Fig. 5a). Currently, the inter-ordinal relationships of the laurasiatherian mammals still remain controversial, as multiple phylogenomic studies gave rise to alternative topologies (Zhou et al. 2012; Foley et al. 2016). The major challenge is ascribed to resolving short internal branches that connect the four key clades (Cetartiodactyla, Perissodactyla, (Carnivora + Pholidota), Chiroptera) that radiated in the late Cretaceous period (Springer et al. 2003). Our result disagrees with the earlier mammal tree (using a supermatrix of 26 gene fragments across 164 mammals) which placed bats as the sister clade to ungulates (((Perissodactyla + Cetartiodactyla), Chiroptera), Carnivora) (Meredith et al. 2011), but supports the recent study (using a supermatrix of 12,931 genes across 48 mammals) that inferred a sister-group relationship between carnivores and ungulates (((Perissodactyla + Cetartiodactyla), Carnivora), Chiroptera) (Jebb et al. 2020). As a perspective of NUMT evolution, our results imply that NUMT presence-and-absence patterns could be an alternative means to infer mammal phylogeny and provide new insights into the resolution of controversial nodes. The power of this approach is expected to increase when more high-quality genomes become available, so that more fixed, orthologous NUMTs are likely to be recovered amongst closely-related species.

Using the ancestral NUMTs on the nodes of the mammal tree and the divergence time, we estimated the NUMT insertion rate for each species (Fig. 5b). However, our insertion rates were overestimated because we were unable to identify NUMTs that originated from post-insertion duplication events. This is due to the intrinsic complexity of NUMT duplication (e.g., tandem duplication; segmental intra- and inter-chromosomal duplication) (Woischnik and Moraes 2002) and NUMT features (e.g., short length; homology to each other; selectively unconstrained) (Leister 2005). Unavailability of chromosome-level genome assemblies has also hindered the identification of these events. We attempted to find tandem duplicated NUMTs that are located within 10 kb windows in genomes, but only a small number of NUMTs meet this criterion (see Methods; Supplemental Table S2). As such, we decided not to consider them when calculating insertion rates. To obtain more precise rates, it is therefore of great importance to analyse multiple chromosome-level genomes across a wide range of species to determine NUMT duplication levels. It was estimated that up to 85% of NUMTs in the human genome originated from *de novo* insertion events (Hazkani-Covo and Graur 2007). Even though evidence for large segmental duplications exists in many species (Bensasson et al. 2003; Triant and DeWoody 2007), this seems to be rare (Hazkani-Covo and Covo 2008). Together with our analyses on tandemly duplicated NUMTs, we hypothesise that NUMT burst in the human, chimpanzee and Tasmanian devil genomes may result from rapid insertion rather than post-insertion duplication.

Although NUMT gain-and-loss patterns over evolutionary time can provide new insights into mammal phylogeny, one should be cautious when using ancestral NUMT trees to infer phylogenetic relationships. These ancestral NUMTs are typically short due to loss in nucleotide sequences over long evolutionary time, and might be subject to different levels of selective constraints in respective species. The erroneous groupings in the trees inferred from Candidates 1 and 2 (Fig. 6) are not unexpected given the short fragments analysed and the fact that several sequence pairs are only overlapping by less than 100 bp. Candidate 3, the NUMT ancestral to Eutheria, was derived from a short fragment of the mtDNA D-loop region which spans a conserved block, and the NUMT sequences have also remained conserved despite ∼100 million years of evolution (only 17 parsimony informative sites across 17 species, Fig. 6). One could combine the information from multiple NUMT alignments to infer relationships, but it is challenging to obtain a consequent number of NUMTs for inferring inter-ordinal relationships in mammals (Fig. 5). Given these limitations, inter-ordinal relationships inferred from ancestral NUMT sequences should be interpreted with caution. That being said, although beyond the scope of our work, more recent NUMTs could be interesting phylogenetic markers to infer intra-ordinal relationships.

To our knowledge, this is the first study that comprehensively explored the characteristics of NUMTs and their evolutionary trajectories in a mammal-wide context. Over the past 200 million years, NUMTs have undergone a fast birth and death in mammalian genomes. Although the reason for their expansions in a few species remains unclear, we revealed that NUMTs are derived from non-random regions in mtDNA, tend to locate in TE-rich and intergenic regions, and unlikely code for functional proteins. Using the new synteny-based method we established, we further demonstrated that NUMT presence-and-absence patterns can provide great insights into mammal evolution, while phylogenies inferred from ancestral NUMTs should be interpreted with caution. As opposed to the traditional alignment-based methods, our novel approach enables NUMT orthology assignment among distantly-related species, providing an alternative means to study phylogeny. This method can potentially be utilised to predict orthology relationships of other fast-evolving, non-coding DNA, such as transposable elements. In the future, a comprehensive taxonomic sampling of species with multiple high-quality individual genomes and a refined genome microsynteny atlas across species will be required to gain a full blueprint of NUMT evolution in mammals. Uniquely, our study broadens the current knowledge on the characteristics of NUMT integrations in mammalian genomes and highlights the merits of NUMT evolution in phylogenetic inference.

## Methods

### Genome sampling

45 published mammalian genomes were used to investigate the evolution of NUMTs in mammals. The list of species comprises Monotremata (n = 1), Marsupialia (n = 4), Afrotheria (n = 4), Xenarthra (n = 1), Lagomorpha (n = 2), Rodentia (n = 5), Scandentia (n = 1), Primates (n = 6), Eulipotyphla (n = 3), Pholidota (n = 1), Carnivora (n = 5), Chiroptera (n = 6), Perissodactyla (n = 2), and Cetartiodactyla (n = 4). These species represent the vast ecological and evolutionary diversity within mammals, representing over ∼200 million years of evolution. We included chicken *Gallus gallus* as a non-mammal outgroup. The quality of genomes was assessed by BUSCO (v4.0) (Waterhouse et al. 2018) and the average genome completeness is 94.0% ± 3.0%, indicative of high quality of these assemblies. The detailed information, including species, genome versions, and genome statistics, is available in Supplemental Table S1.

### Optimization of NUMT identification pipelines

For each species, we employed a local BLAST approach (Altschul et al. 1990) to identify NUMT insertions by querying its nuclear genome using the corresponding complete mitochondrial genome sequences. Where the mitogenome of the same species was not available, that of its closely-related species was used (see Supplemental Table S1). To facilitate BLAST search, circular mitogenomes were presented as linear sequences that begin with tRNA-Phe and end with D-loop.

There are currently no standard pipelines and criteria to define NUMT insertions. To best profile NUMTs in mammalian genomes, we tested the default BLASTN (v2.9.0) and discontinuous mega-BLAST (v1.6.0; more sensitive to detect divergent sequences) with a combination of three key parameters (*-template_length*; *-template_type, -word_size*) using the human genome (hg38) as an example (Supplemental Table S8). We applied a stringent E-value threshold (10^−6^) and a minimum HSP (High-scoring Segment Pair) length (30 bp) to avoid potential false-positives that were derived from assembly errors or non-mitochondrial origin. The BLASTN with default parameter settings only produced 186 HSPs, with many old NUMTs (less than 80% identity to their corresponding mtDNA) undetected. In contrast, discontinuous mega-BLAST generally reported more NUMTs, and different combinations of parameters yielded similar HSP numbers (839 ± 8) which are higher than most of previous studies reported (Hazkani-Covo and Graur 2007; Hazkani-Covo 2009) (Supplemental Table S8). Based on the sensitivity, we chose discontinuous mega-BLAST with the following parameters: *-template_length 18, -template_type optimal* and *-word_size 11* as the optimal approach.

### Identification and characteristics of NUMTs in 46 genomes

Using the optimal method, we predicted NUMTs in all 46 genomes. For each species, the raw BLAST HSPs were merged if they meet the following conditions: 1) there is a less than 10 bp gap in the nuclear genome between two adjacent HSPs that have continuous mitogenome coordinates correspondingly; 2) a single NUMT that traverses the D-loop region are split into two HSPs due to the boundary created by linearization of circular mtDNA. We further removed the HSPs located in very short contigs (< 20 kb) as they were likely the result of mtDNA contamination or assembly errors. Subsequently, for each species we assigned continuous numbers (e.g., Hsap_numt_1) to the processed HSPs according to their nuclear genomic coordinates (Supplemental Table S3).

To explore the link between NUMT insertion and nuclear genome, we performed correlation analyses between some NUMT characteristics (number; accumulative length) and genome statistics (genome size; scaffold number; transposable element content) using Spearman’s correlation tests. Significances of the tests were further corrected by phylogeny using the *phytools* R package (Revell 2012). The time-calibrated phylogenetic tree was obtained from the recent published mammal phylogeny (Jebb et al. 2020). Next, we examined if there are any ‘hotspots’ or overabundance in mtDNA from which NUMTs were derived. To do this, we obtained the mitochondrial cross coverage of NUMTs across species using *genomecov* in the BEDTools suite (v2.30.0) (Quinlan and Hall 2010). We firstly scanned the coverage of mtDNA with a 50 bp sliding window and calculated the median coverage per window for each species. Then, we conducted comparisons of the coverage between all possible windows across species using Mann-Whitney *U* tests. Due to the disparity on mtDNA length across species, we only analysed the first 320 windows representing 16,000 bp in mtDNA (1^st^ ∼ 16,000^th^ bp), starting with the gene tRNA-Phe. To further confirm our results, we simulated a null distribution of mtDNA coverage by randomly reshuffling mitogenome coordinates of NUMTs for each species, performed pairwise comparisons of the coverage between all windows as mentioned above, and repeated these analyses 1,000 times. For the coordinates, we only randomly picked a start position for each NUMT (between 1 ∼ 16,000) and used the length of observed NUMTs to calculate the end coordinates. Hence, the length distribution of NUMTs was identical between the observed and simulated datasets. In order to take mtDNA circularity into account, coordinates greater than 16,000 (e.g. 16,200 ∼ 16,450) were subtracted 16,000 (e.g. leading to 200 ∼ 450). Finally, the number of significant tests obtained from the observed dataset was compared to the distribution obtained from the 1,000 simulated datasets (i.e. null distribution if coverage was homogenous).

### NUMT phylogenetic trees

To explore the NUMT evolution in mammals, we built phylogenetic trees using NUMT sequences across 45 species. We obtained all NUMT sequences that are mapped to two mitochondrial genome loci, Cytochrome b (*CYTB*) and NADH dehydrogenase 1 (*ND1*), which are commonly used as genetic markers in phylogenetic studies. For each locus, we only selected the mapped NUMT sequences that are equal or over 500 bp (188 sequences for *CYTB* and 215 sequences for *ND1*), and aligned them together with mtDNA of 45 species using MAFFT (Katoh and Standley 2013). The appropriate nucleotide substitution models (GTR+F+G4 for CYTB and K3Pu+F+R6 for ND1) were selected based on the Bayesian information criterion (BIC) using ModelFinder (Kalyaanamoorthy et al. 2017) (Supplemental Table S9). Next, we inferred the maximum-likelihood (ML) tree using the partition model in IQ-TREE (Nguyen et al. 2015; Chernomor et al. 2016) for each locus. To search for the best-scoring ML, we performed ultrafast bootstrap (UFBoot) (Hoang et al. 2018) with 1,000 bootstraps and 1,000 topology replicates. To verify the robustness of the ML trees, the branch supports were evaluated using SH-like approximate likelihood ratio test (SH-aLRT) (Guindon et al. 2010) and a Bayesian-like transformation of aLRT (aBayes) (Anisimova et al. 2011). SH-aLRT was performed with 1,000 replicates. The ML, SH-aLRT, and aBayes analyses were performed using W-IQ-TREE (Trifinopoulos et al. 2016). The trees were rooted by Monotremata.

### Assembly of NUMT blocks

To facilitate the downstream analyses, for each species the adjacent NUMTs with a nuclear genomic distance less than 2 kb were assembled as a single NUMT block, regardless of their orientations in the mitogenome (Supplemental Table S4). NUMT blocks are considered complex if they are comprised of three or more individual NUMTs. To demonstrate the reliability of predicted large NUMT blocks, as an example, we examined their nuclear genomic loci using our published PacBio raw reads for *Molossus molossus* bat genome assembly (Supplemental Fig. S2c), given that *M. molossus* possesses one of the largest NUMT blocks amongst six bat species we studied.

### Analyses of NUMT insertion ‘hotspots’ in nuclear genomes

To understand if there are any ‘hotspots’ or preference in nuclear genomic regions in which NUMT insertions occurred, we firstly investigated the TE content in the flanking regions of NUMTs/NUMT blocks in genomes. For each species we extracted 5 kb flanking sequences, both upstream and downstream, of each NUMT/NUMT block using *getfasta* in the BEDTools suite (v2.30.0) (Quinlan and Hall 2010). Their TE contents were estimated using RepeatMasker (v4.1.2) (Smit 2013-2015) with a window size of 500 bp. We compared the average TE content in 5 kb flanking regions of all the NUMTs/NUMT blocks at both ends (20 windows) against the average genome TE content in each species using Mann-Whitney *U* tests. Using this method we also compared the TE content between different windows in a pairwise manner. It is noted that TE content may be underestimated because the TE database is currently biased for only a few model species such as human and mouse (Smit 2013-2015). Because TE were compared across windows of NUMT flanking regions within species, this bias is unlikely to affect our conclusions. To further explore if newly-inserted NUMTs are located in proximity to TE, for each species we performed correlation analyses between NUMT/mtDNA sequence identity and the distance of NUMTs to their closest TE (averaged by both ends) using Spearman’s correlation tests. Owing to the heterogeneity in sequence identity of individual NUMTs, NUMT blocks were excluded from this analysis.

Next, we investigated if NUMTs are prone to insert into intronic or intergenic regions. To achieve this we obtained the high-quality genome annotation files published along with the genomes from the National Center for Biotechnology Information (NCBI). We did not include the following species: *Rattus norvegicus, Sciurus vulgaris, Saimiri boliviensis, Manis javanica, Gymnobelideus leadbeateri, Ceratotherium simum, Trichechus manatus, Tupaia belangeri* and *Vombatus ursinus* in the analysis, because the genome annotation files of these species were not available or of poor quality (Supplemental Table S1). For the remaining 36 species, non-coding gene annotations were removed from their genome annotation files. Intronic and intergenic NUMTs were determined by merging their genomic coordinates with protein-coding gene coordinates using *merge* in the BEDTools suite (v2.30.0) (Quinlan and Hall 2010). However, there is a caveat that intergenic NUMTs close to protein-coding genes might be located in gene untranslated regions (UTRs). This is because UTRs are typically not well annotated in most of mammalian genomes.

### Functional predictions of NUMTs

To ascertain if NUMTs are expressed and functional, we obtained and analysed publicly available RNA-Seq data of four tissue types (brain, kidney, liver and heart) from five species (human, naked mole-rat, cow, dog and platypus) (Supplemental Table S10). Because NUMT sequences could be very similar to their corresponding mtDNA sequences depending on their insertion time, we used the stringent criteria to determine if a NUMT is expressed. NUMTs are considered expressed if 1) at least two RNA-Seq reads support the junctions between NUMTs and their flanking nuclear genomic regions with at least 5 bp overhangs, and 2) the coverage of NUMTs by RNA-Seq reads is >70%. To achieve this, for each species we extracted the sequence of each NUMT/NUMT block together with their 200 bp upstream and downstream flanking sequences as references using the same method mentioned above. Prior to NUMT quantification, adaptors and low-quality regions (base score < Q25) in raw RNA-Seq reads were filtered using cutadapt (v3.5) (Martin 2011). We then mapped the clean reads from each sample to the corresponding references using HISAT2 (v2.2.1) (Kim et al. 2015). NUMT expression was analysed using Samtools (v1.13) (Li et al. 2009) and the BEDTools suite (v2.30.0) (Quinlan and Hall 2010), and was further visualised in the genome browser IGV (v2.14.1) (Robinson et al. 2011). Next, we explored if NUMT sequences have the potential to be translated into proteins. To achieve this, we investigated all 17,732 individual NUMTs across 45 mammals. We firstly identified NUMTs that contain the entire regions of any mtDNA protein-coding genes, and analysed their open reading frames (ORFs) using Geneious (v11.0.5) (https://www.geneious.com). ORFs were then translated into proteins based on both nuclear and mitochondrial genetic codes (Supplemental Table S5).

### A novel method to determine NUMT orthology between distant-related species

We initially attempted to align NUMT sequences along with 1 kb flanking sequences at each end in each species using Clustalw (v2.1) (Thompson et al. 1994). With the exception of the closely-related human and chimpanzee which diverged only ∼5 Myr ago, NUMTs with flanking regions were poorly aligned amongst the remaining species (data not shown), and thus, the results were not conclusive. As such, traditional alignment-based methods are not feasible to infer orthologous NUMTs between distant-related species.

To address this problem, we established an innovative and practical approach that utilises genome microsynteny to identify orthologous NUMTs within mammalian clades. We used protein-coding genes in a conserved genomic synteny block as anchors to infer orthologous NUMTs/NUMT blocks among species. We regarded two NUMTs/NUMT blocks from respective species as orthologous, if 1) they are located in the same synteny block within a distance of 6 protein-coding genes (3 genes upstream and 3 genes downstream of a NUMT/NUMT block), and 2) their sequences overlap with each other by at least 50% (Supplemental Fig. S8a). With these criteria, we evaluated the probability (error rate) that two NUMTs/NUMT blocks from respective species were assigned as orthologs by chance. Due to the complexity of NUMT insertions (e.g. different NUMT lengths; complicated NUMTs) and different characteristics of genomes (e.g. different numbers of protein-coding genes), we employed a simplified formula to estimate the error rate of NUMT orthology assignment for each pair of species (Supplemental Fig. S8b). Suppose that the average number of protein-coding genes in mammalian genomes is 20,000, that the average length of NUMTs is 200 bp, and that the average size of mammalian mitochondrial genomes is 16,600 bp. The error rate was calculated at 3.03 × 10^−6^ (see Supplemental Fig. S8b for the explanation). The mathematical error expectation (*E*) of orthology assignment was estimated by multiplying the error rate (3.03 × 10^−6^) by the total number of all possible NUMT pairs (*N*_*a*_ × *N*_*b*_) between two species (Supplemental Fig. S8c). *N*_*a*_ and *N*_*b*_ stand for the number of NUMTs/NUMT blocks in species A and B, respectively. With the exception of the comparisons between *S. harrisii* and the other species, the estimated error expectations (*E*) amongst the remaining species are far below 1 (Supplemental Fig. S8c). These results imply that our novel approach is accurate and feasible to assign NUMT orthology between distantly-related species within mammals.

### Determination of orthologous NUMTs across mammals

Using this method, we predicted orthologous NUMTs/NUMT blocks between species by leveraging 630 pairwise comparisons of genome-wide microsynteny across 36 mammals in which high-quality gene annotation files are available. For each species, we integrated the predicted NUMT annotations with the protein-coding gene annotations using *merge* in the BEDTools suite (v2.30.0) (Quinlan and Hall 2010), and assigned orthologous NUMTs/NUMT blocks between two species on the basis of the above criteria. Each predicted pair was manually inspected. The NUMT orthology between two species was visualised using the R package *circlize* (v0.4.15) (Gu et al. 2014), and the NUMT orthology networks were established using the R package *UpSetR* (v1.4.0) (Conway et al. 2017). To facilitate data visualisation and interpretation, we categorised these 36 species into eight clades with a balanced species number per clade. These defined clades include Clade 1 (Monotremata + Marsupialia, n = 3), Clade 2 (Afrotheria + Xenarthra, n = 4), Clade 3 (Primates, n = 5), Clade 4 (Rodentia + Lagomorpha, n = 5), Clade 5 (Eulipotyphla, n =3), Clade 6 (Carnivora, n = 5), Clade 7 (Perissodactyla + Cetartiodactyla, n = 5), and Clade 8 (Chiroptera, n = 6). It is noteworthy that we predicted 6 orthologous NUMTs/NUMT blocks between *O. anatinus* and *S. harrisii* (Clade 1; Supplemental Fig. S9a). Given their long divergence time and our estimation of the error expectation (*E* = 2.99; Supplemental Fig. S8c), it is likely that these predictions are false-positives mainly because *S. harrisii* has the largest number of NUMTs (Fig. 1a).

### Ancestral orthologous NUMTs on the phylogenetic tree and NUMT insertion rates

Because most NUMTs are non-functional and under limited selective constraints, we employed a simple coalescent method to infer ancestral NUMTs/NUMT blocks on the nodes of the given phylogenetic tree (Jebb et al. 2020) using the phylogenetic patterns of NUMT presence-and-absence. A NUMT/NUMT block is regarded as ancestral on the node if it is identified as orthologous across species in both bifurcating clades to which the node branches. Next, we estimated the NUMT insertion rate as described in (Hazkani-Covo 2009). For each species we obtained the number of individual NUMTs that do not have orthologs in its most closely-related species or monophyletic group. The NUMT insertion rate (number of insertions per 1 million years) for each species was estimated by dividing the number of individual, non-orthologous NUMTs by the divergence time. It is noteworthy that it is particularly challenging to identify duplications of pre-existing NUMTs due to the complexity of these events and lack of chromosome-level genome assemblies. We attempted to identify tandem duplicated NUMTs which are located within 10 kb windows, have similar start or end mitogenome coordinates (± 10 bp), and overlap with each other by at least 50% for each species. These duplicated NUMTs were further verified by aligning them along with 1 kb flanking sequences at both ends using Clustalw (v2.1) (Thompson et al. 1994). Due to the scarcity of this case observed across mammals (Supplemental Table S1), we decided not to consider duplication events when calculating the insertion rate.

### Alignments of ancient ancestral orthologous NUMTs across species

Next, we visualised the genomic positions of all the seven ancestral NUMTs/NUMT blocks in genomic microsynteny across species (Fig. 6; Supplemental Fig. S10), and constructed three phylogenetic trees for the individual ancestral NUMTs (Candidates 1-3) respectively. The sequences of Candidates 1-3 were aligned and trimmed to the shortest common length using Gblocks (v0.91b) (Castresana 2000). The phylogenetic trees were inferred and verified as extensively described above (see NUMT phylogenetic trees). For Candidates 1-3, the alignment lengths are 228 bp (17 sequences), 342 bp (15 sequences) and 253 bp (17 sequences), respectively. The best-fit models are HKY+F+I, TPM3+F and TPM2u+F, respectively.

### Statistical analyses

The statistical analyses used in this study, including Man-Whitney *U* tests, Kolmogorov-Smirnov tests, Spearman’s correlation tests and Chi-square tests were performed in R (v4.1.1) (Team 2014). *P*-values were corrected by multiple tests using false discovery rate (FDR) where applicable. Statistical tests with corrected *P* < 0.05 were considered significant unless specifically defined.

## Data access

The publicly available genome, mitogenome and RNA-Seq data used in this study are documented in the Supplemental Tables 1 and 10. The intermediate data supporting the conclusions can be available at the GitHub page (https://github.com/huangzixia/NUMT_evolution_in_mammals).

## Acknowledgements

We thank Emmanuel Douzery from University of Montpellier for discussions on phylogenetic analyses. We also acknowledge the UCD Sonic High Performance Computing for the provision of computational facilities and support. This study is supported by the Irish Research Council Laureate Bursary Grant (No.74725) and the UCD seed funding (No.68674) to Z.H., the Irish Research Council Laureate Award IRCLA/2017/58 and Science Foundation Ireland Future Frontiers 19/FFP/6790 awarded to E.C.T., and the Junior Chair from Institut Universitaire de France awarded to S.J.P.

## Author contributions

Z.H. developed the novel approach for NUMT orthology assignment across species and devised all the analyses. Z.H., M.U., S.J.P., S.P., M.P. and S.C. performed the analyses. Z.H., M.U., S.J.P., E.C.T., W.H. and E.W.M. interpreted the results. Z.H. and M.U. are responsible for the figures and tables presented throughout. Z.H. wrote the first draft, with input from all the authors.

