## Supplemental Figures S1-S10, Tables S8-S10 for "Comparative genome microsynteny illuminates the fast evolution of nuclear mitochondrial segments (NUMTs) in mammals"

**This pdf file contains:**

**Supplemental Figures S1 – S10**

**Supplemental Tables S8 – S10**

***Supplemental Tables S1 – S7 are provided as separate Excel files (.xlsx)**

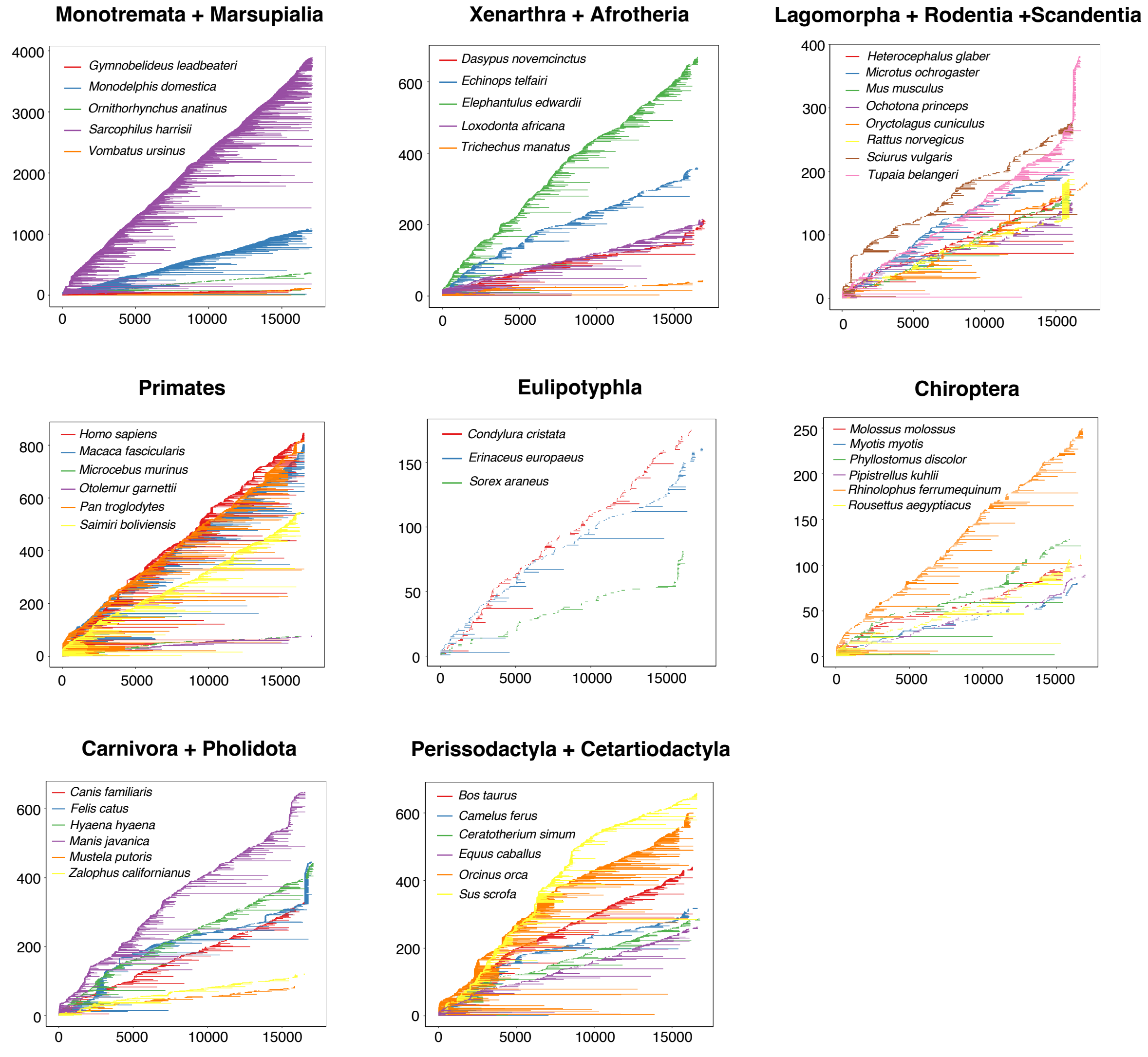


**Supplemental Fig. S1: Coordinates in the mitochondrial genome matching NUMTs (hsps) in each of 45 mammalian genomes**. The *x*-axis represents the nucleotide positions in mitochondrial genomes and the *y*-axis represents the number of NUMTs (hsps). Because mitogenomes are circular and were thus linearized, the map begins with the first tRNA, tRNA-Phe, immediately before 12s RNA. To better present the data, NUMT coordinates in 45 genomes were visualised in 8 panels.

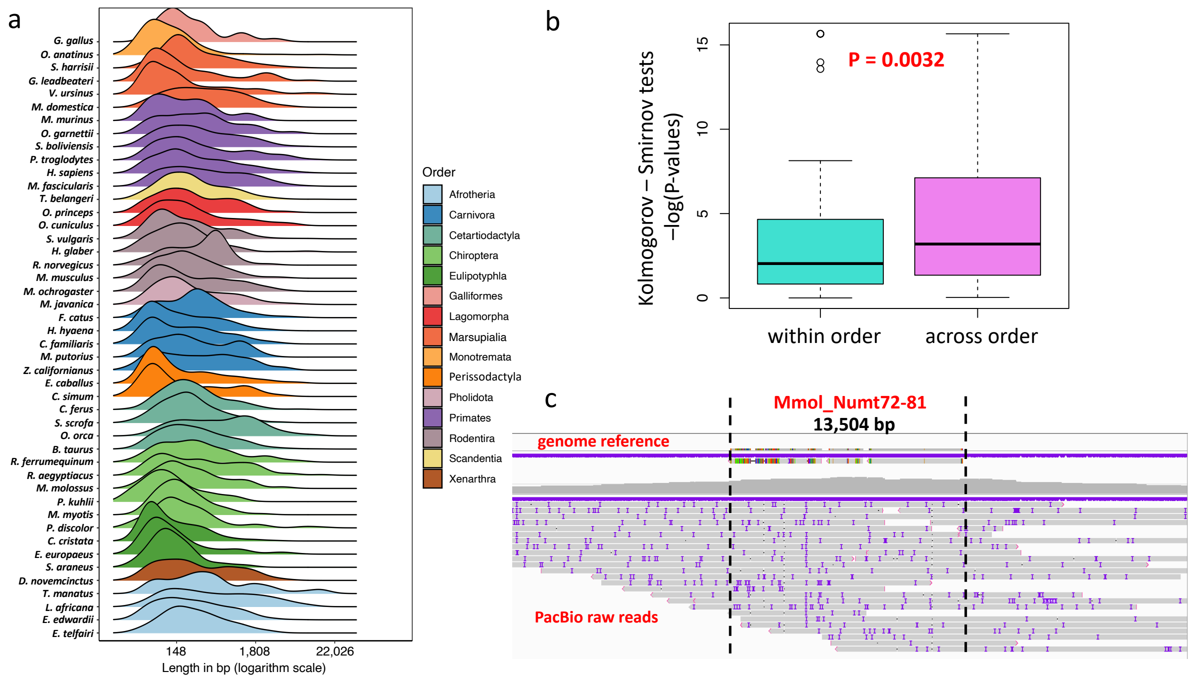


**Supplemental Fig. S2: NUMT length in mammalian genomes. a)** NUMT length distribution across 45 mammalian genomes plus *G. gallus* as the outgroup. The scale of the *x*-axis (NUMT length) was log *e* transformed. **b)** Comparisons of NUMT length distributions across species between and within orders. Pairwise comparisons of NUMT length distributions between species were conducted using Kolmogorov-Smirnov tests. *P*-values obtained from the tests of within-order and across-order comparisons were ($-\log_{10}$) transformed, and further compared using Mann-Whitney *U* tests. **c)** An example of authentication of long, complicated NUMTs using PacBio reads. One large NUMT block (Mmol_Numt72-81, 13,504 bp in total) in the *M. molossus* genome shows that a number of PacBio raw reads used for genome assembly support the junctions between the NUMT block and flanking genomic sequences. The two dashed lines indicate the boundaries between the NUMT block and its up- and down-stream flanking genomic regions. The reference genome and PacBio raw reads are indicated on the graph accordingly.

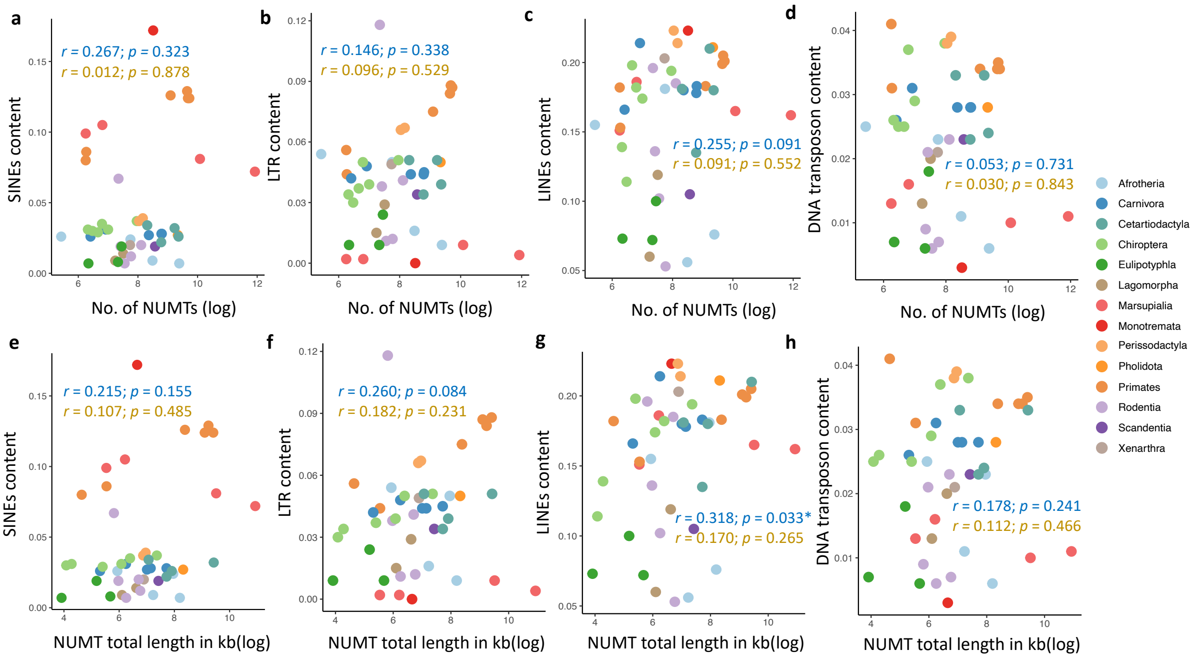


**Supplemental Fig. S3: Correlation between NUMT features and genome transposable element (TE) content.** TEs that were investigated include short interspersed nuclear elements (SINEs), long interspersed nuclear elements (LINEs), long terminal repeats (LTRs), and DNA transposons. **a-d)** Scatterplots showing the correlation between NUMT (hsp) number, and genome SINEs, LINEs, LTRs and DNA transposon content, respectively. **e-h)** Scatterplots showing the correlation between total NUMT length, and genome SINEs, LINEs, LTRs and DNA transposon content, respectively. Correlation coefficients ($r$) and $P$-values were computed using Spearman's correlation tests. In the scatterplots, coefficients ($r$) and $P$-values in blue and gold indicate the values before and after phylogeny correction (*$0.01 < P < 0.05$). The colour code indicates the species from the same order.

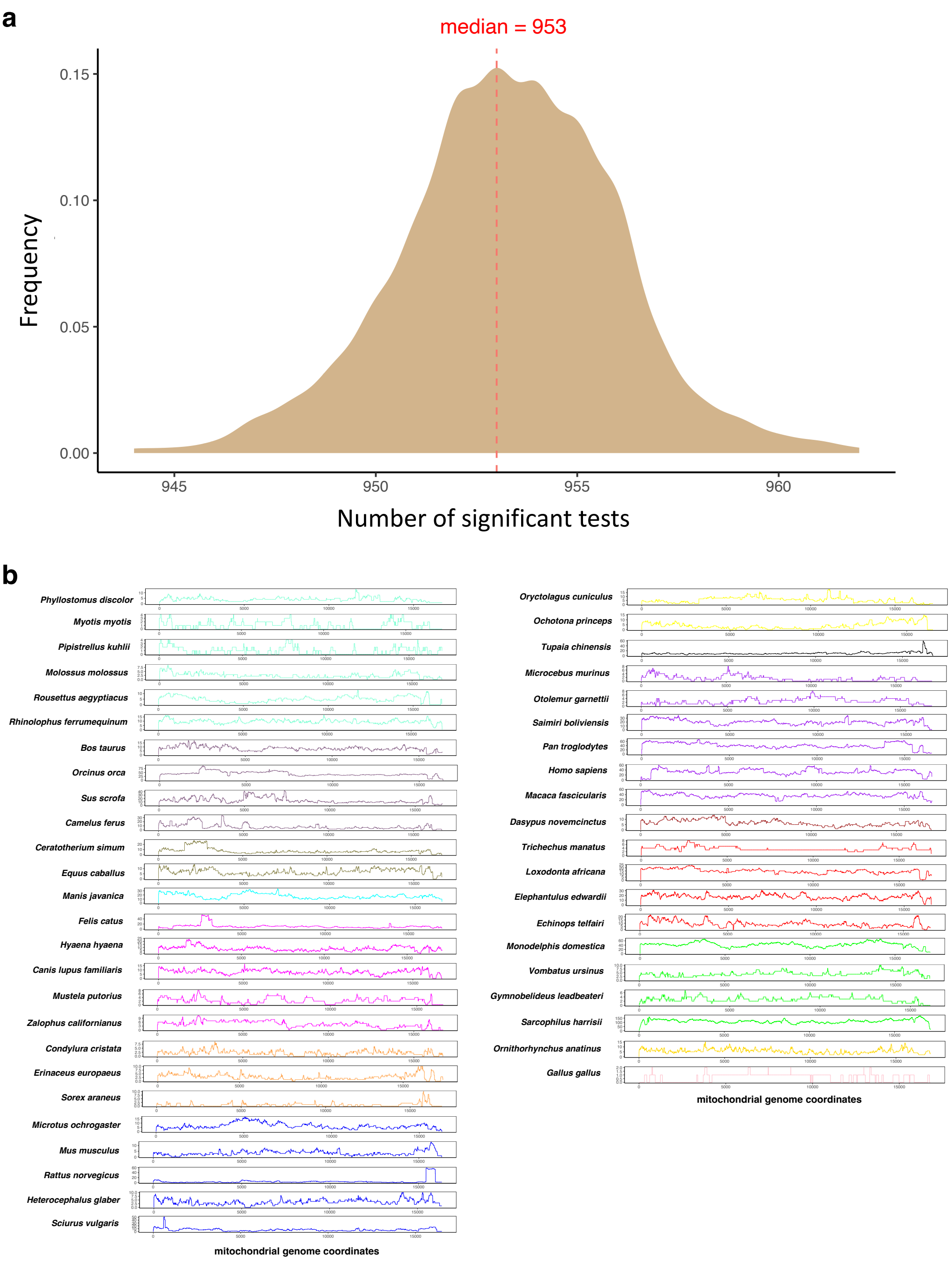


**Supplemental Fig. S4: Coverage of NUMTs in mtDNA. a)** Distribution of significant tests between NUMT coverage of all possible windows from 1,000 simulated datasets. Out of 51,040 tests, the median number of the significant tests is 953. **b)** The coverage of NUMTs (hsps) in the linearised mtDNA in 45 mammalian genomes and the *G. gallus* genome. The *x*-axis indicates the position in the mitogenome and the *y*-axis indicates the number of NUMTs which overlap with the mitogenome at each position. The linearised mtDNA begins with tRNA-Phe and ends with D-loop. The colour code indicates the species from the same orders.

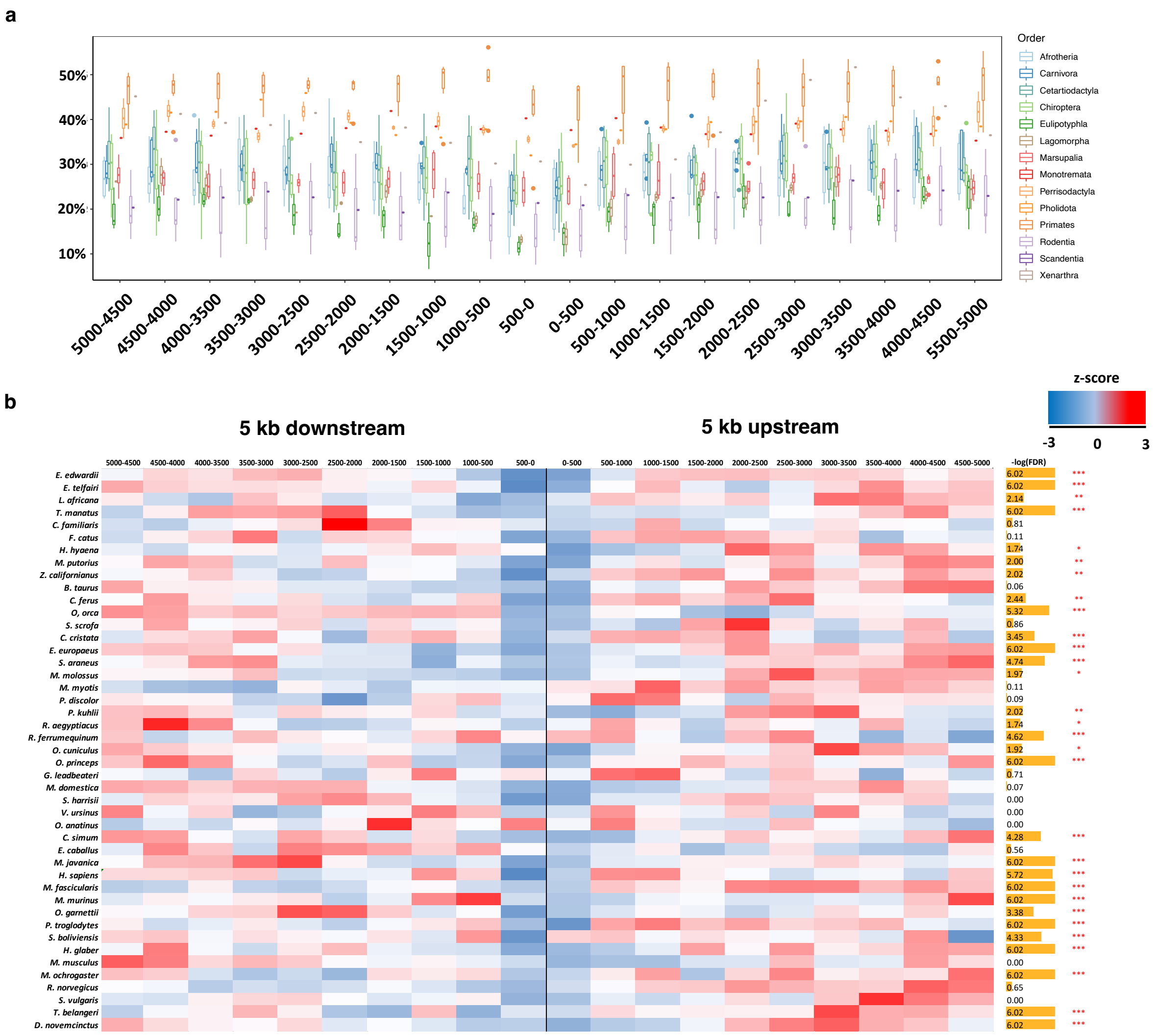


**Supplemental Fig. S5: Transposable element (TE) content in the 5kb up- and down-stream flanking regions of NUMTs/NUMT blocks. a)** The non-normalised average TE contents in the 5kb up- and down-stream flanking regions of NUMTs/NUMT blocks with a window size of 500 bp across 45 mammals. In the boxplot the species were grouped into the respective orders. **b)** The heatmap showing the average TE content in the 5kb up- and down-stream flanking regions of NUMTs/NUMT blocks with a window size of 500 bp for each species. The average TE content of 20 windows per species were normalized to Z-scores. Blue: low TE content; white: median TE content; red: high TE content. For each species, the average TE contents of 20 windows were compared to the genome average TE content using Mann-Whitney $U$ tests (***$P < 0.001$; **$0.001 < P < 0.01$; *$0.01 < P < 0.05$).

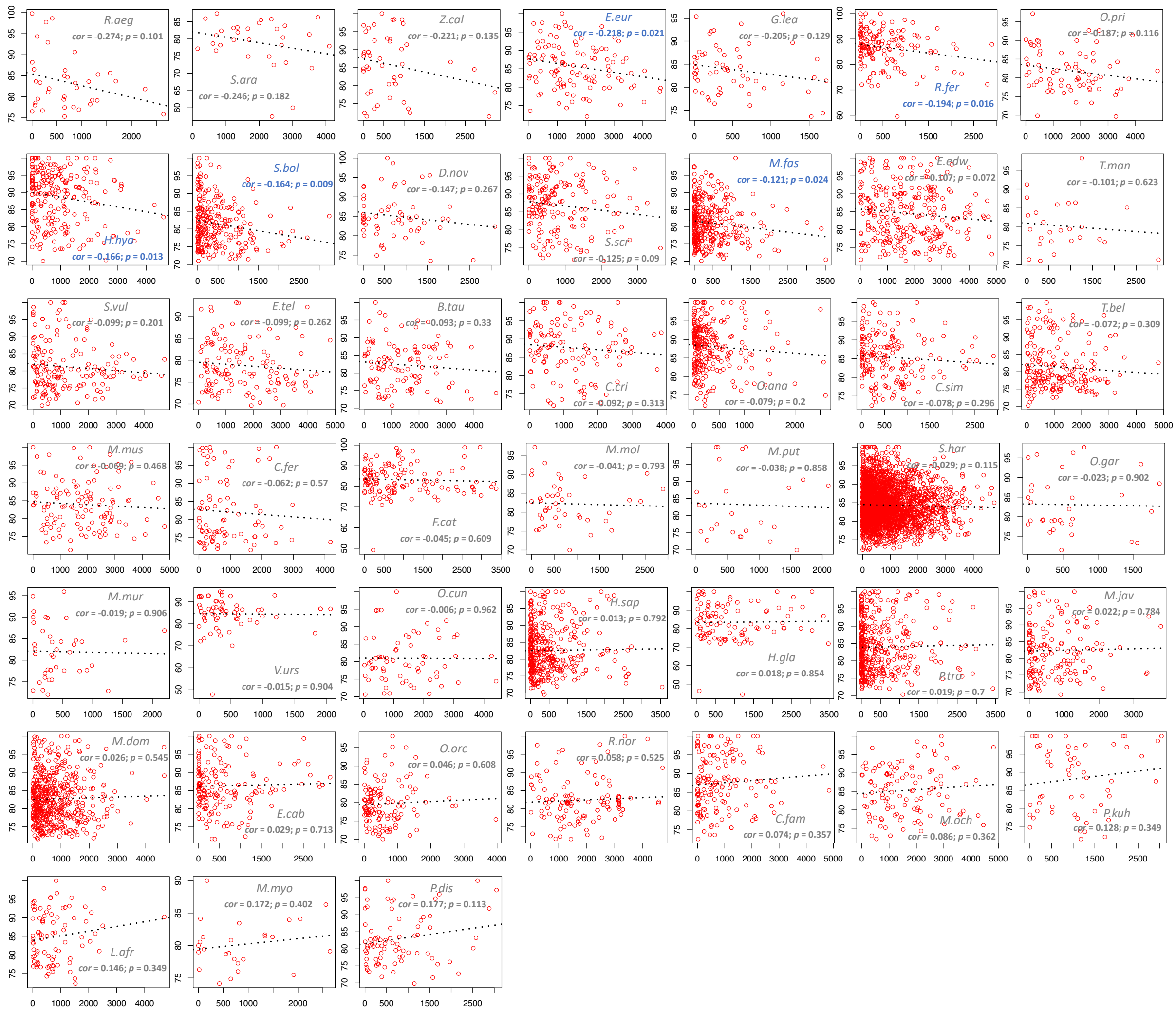

**Supplemental Fig. S6: Correlation between NUMT distance to the closest TE and the sequence identity to their corresponding mtDNA regions across 45 mammals.** The *x*-axis represents the distance (bp) between NUMTs and their closest TE, and the *y*-axis represents the sequence identity (%) between NUMTs and their corresponding mtDNA regions. The correlation coefficient was calculated using a Spearman correlation test for each species. The regression line was inferred using a generalized linear model. Five species (in blue) exhibit a significant negative correlation between NUMT distance to the closest TE and NUMT sequence identity to the corresponding mtDNA regions.

a

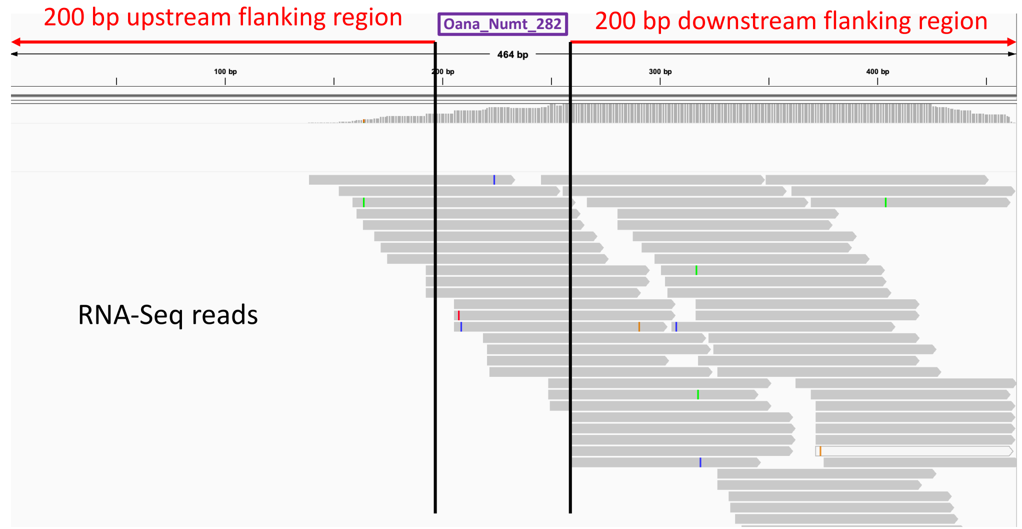


b

| | Brain | Kidney | Liver | Heart |
|---|---|---|---|---|
| **Platypus** | 5.26% | 5.76% | 3.51% | 3.51% |
| **Naked mole-rat** | 0 | 0.81% | 1.63% | 0.81% |
| **Human** | 0.39% | 0 | 0 | 0 |
| **Dog** | 0.61% | 1.82% | 1.21% | 1.21% |
| **Cow** | 0 | 0 | 0 | 0.61% |

c

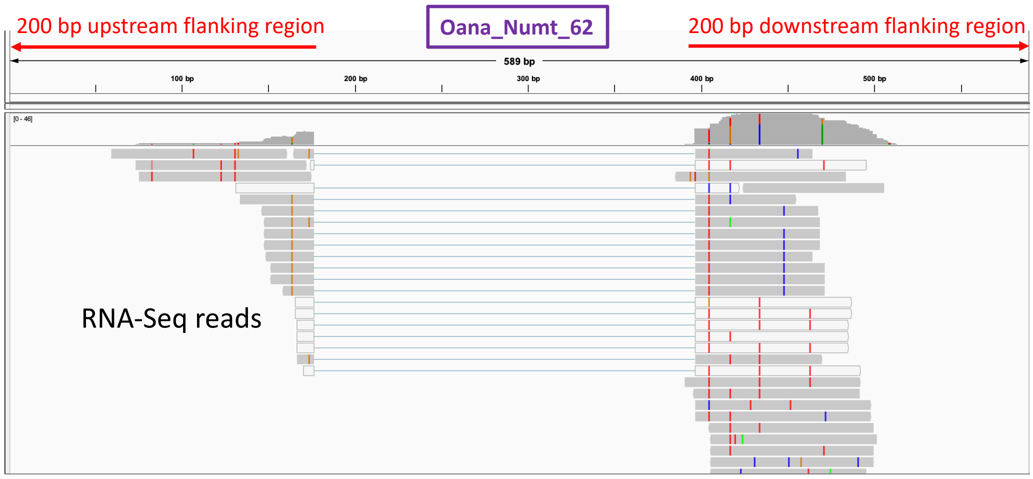


**Supplemental Fig. S7: NUMT expression analyses. a)** An example of expressed NUMTs. The expression of Oana_numt282 in platypus is supported by a number of RNA-Seq reads mapped to the boundaries between the NUMT and its flanking regions at both ends. **b)** The percentage of NUMTs/NUMT blocks expressed in 4 tissue types across 5 mammals. We averaged the percentage of NUMTs/NUMT blocks across samples from the same tissue type for each species. **c)** An example of polymorphic NUMTs revealed by RNA-Seq reads. Mapped RNA-Seq reads were spliced at the Oana_numt62 locus in the platypus genome. This is due to the fact that the sample sources for the genome and RNA-Seq sequencing came from different individuals.

**a**

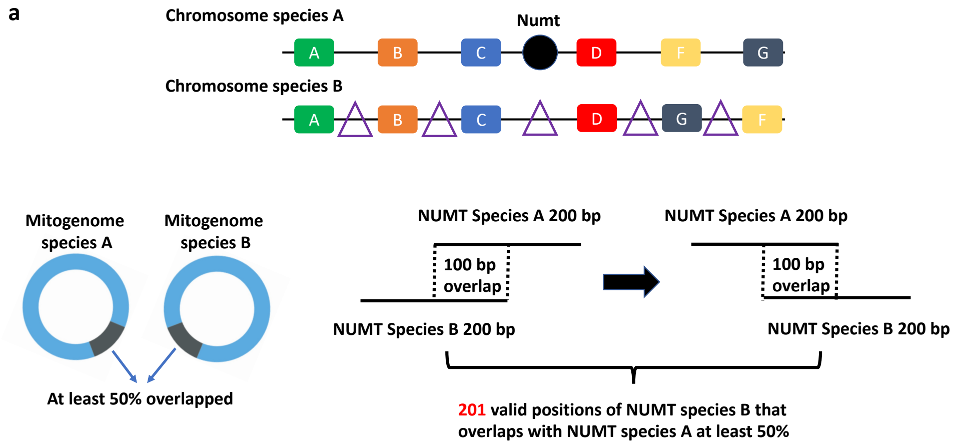


**b** Suppose:

1) the average number of protein-coding genes in mammalian genomes is **20,000** (***N***)
2) The median length of Numts in mammals is **200 bp**. There are **201** valid positions ($N_{vp}$) if two orthologous Numts (species A and B) overlap at least 50% of their sequences.
3) The median length of mitogenomes in mammals is **16,600 bp** (***NA, NB***) and the number of possible positions with which Numts can start is **16,600** (***N***).

$$\text{Probability (P)} = \left\{ \frac{1}{N+1} \times 5 \right\} \times \left\{ \frac{1}{NA} \times \frac{1}{NB} \times N \times N_{vp} \right\}$$

$$P = \left\{ \frac{1}{20000+1} \times 5 \right\} \times \left\{ \frac{1}{16600} \times \frac{1}{16600} \times 16600 \times 201 \right\} = 3.03 \times 10^{-6}$$

**c**

| | Mmol | Mmyo | Pkuh | Pdis | Raeg | Rfer | Ecab | Sscr | Cfer | Btau | Oorc | Fcat | Cfam | Mput | Hhya | Zcal | Eeur | Ccri |
|---|---|---|---|---|---|---|---|---|---|---|---|---|---|---|---|---|---|---|
| Mmol | | 0.01 | 0.01 | 0.02 | 0.01 | 0.03 | 0.03 | 0.05 | 0.03 | 0.03 | 0.04 | 0.03 | 0.03 | 0.01 | 0.05 | 0.01 | 0.01 | 0.03 |
| Mmyo | | | 0.01 | 0.01 | 0.01 | 0.02 | 0.03 | 0.03 | 0.02 | 0.02 | 0.03 | 0.03 | 0.03 | 0.01 | 0.03 | 0.01 | 0.01 | 0.01 |
| Pkuh | | | | 0.02 | 0.01 | 0.03 | 0.04 | 0.05 | 0.03 | 0.03 | 0.05 | 0.04 | 0.04 | 0.01 | 0.06 | 0.01 | 0.01 | 0.03 |
| Pdis | | | | | 0.01 | 0.05 | 0.06 | 0.08 | 0.04 | 0.05 | 0.07 | 0.05 | 0.06 | 0.01 | 0.09 | 0.02 | 0.01 | 0.04 |
| Raeg | | | | | | 0.03 | 0.03 | 0.05 | 0.03 | 0.03 | 0.04 | 0.03 | 0.03 | 0.01 | 0.05 | 0.01 | 0.01 | 0.03 |
| Rfer | | | | | | | 0.11 | 0.15 | 0.08 | 0.09 | 0.13 | 0.11 | 0.11 | 0.02 | 0.16 | 0.03 | 0.01 | 0.07 |
| Ecab | | | | | | | | 0.16 | 0.09 | 0.10 | 0.15 | 0.12 | 0.13 | 0.03 | 0.18 | 0.04 | 0.02 | 0.09 |
| Sscr | | | | | | | | | 0.11 | 0.13 | 0.19 | 0.16 | 0.17 | 0.03 | 0.23 | 0.05 | 0.03 | 0.11 |
| Cfer | | | | | | | | | | 0.07 | 0.11 | 0.09 | 0.09 | 0.02 | 0.13 | 0.03 | 0.01 | 0.06 |
| Btau | | | | | | | | | | | 0.12 | 0.09 | 0.10 | 0.02 | 0.14 | 0.03 | 0.01 | 0.07 |
| Oorc | | | | | | | | | | | | 0.14 | 0.15 | 0.03 | 0.21 | 0.05 | 0.02 | 0.10 |
| Fcat | | | | | | | | | | | | | 0.12 | 0.03 | 0.17 | 0.04 | 0.02 | 0.08 |
| Cfam | | | | | | | | | | | | | | 0.03 | 0.18 | 0.04 | 0.02 | 0.09 |
| Mput | | | | | | | | | | | | | | | 0.04 | 0.01 | 0.01 | 0.02 |
| Hhya | | | | | | | | | | | | | | | | 0.05 | 0.03 | 0.12 |
| Zcal | | | | | | | | | | | | | | | | | 0.01 | 0.03 |
| Eeur | | | | | | | | | | | | | | | | | | 0.01 |

| | Sara | Mmus | Moch | Hgla | Ocun | Opri | Hsap | Ptro | Mfas | Mmur | Ogar | Dnov | Lafr | Etel | Eedw | Mdom | Oana | Shar |
|---|---|---|---|---|---|---|---|---|---|---|---|---|---|---|---|---|---|---|
| Mmol | 0.01 | 0.02 | 0.03 | 0.02 | 0.02 | 0.02 | 0.09 | 0.09 | 0.09 | 0.01 | 0.01 | 0.02 | 0.02 | 0.03 | 0.07 | 0.13 | 0.05 | 0.58 |
| Mmyo | 0.01 | 0.01 | 0.02 | 0.01 | 0.01 | 0.01 | 0.06 | 0.07 | 0.06 | 0.01 | 0.01 | 0.01 | 0.01 | 0.03 | 0.05 | 0.09 | 0.03 | 0.40 |
| Pkuh | 0.01 | 0.03 | 0.03 | 0.03 | 0.02 | 0.02 | 0.10 | 0.11 | 0.09 | 0.01 | 0.01 | 0.02 | 0.02 | 0.04 | 0.09 | 0.14 | 0.06 | 0.65 |
| Pdis | 0.01 | 0.04 | 0.05 | 0.03 | 0.03 | 0.03 | 0.15 | 0.16 | 0.14 | 0.01 | 0.01 | 0.03 | 0.03 | 0.05 | 0.13 | 0.21 | 0.09 | 0.97 |
| Raeg | 0.01 | 0.02 | 0.03 | 0.02 | 0.02 | 0.01 | 0.09 | 0.09 | 0.09 | 0.01 | 0.01 | 0.02 | 0.02 | 0.03 | 0.07 | 0.12 | 0.05 | 0.57 |
| Rfer | 0.03 | 0.07 | 0.08 | 0.07 | 0.06 | 0.05 | 0.28 | 0.29 | 0.27 | 0.03 | 0.03 | 0.05 | 0.07 | 0.11 | 0.23 | 0.39 | 0.16 | 1.79 |
| Ecab | 0.03 | 0.08 | 0.09 | 0.07 | 0.07 | 0.06 | 0.31 | 0.33 | 0.30 | 0.03 | 0.03 | 0.06 | 0.07 | 0.12 | 0.26 | 0.43 | 0.18 | 2.01 |
| Sscr | 0.04 | 0.11 | 0.13 | 0.10 | 0.09 | 0.08 | 0.42 | 0.45 | 0.39 | 0.04 | 0.04 | 0.08 | 0.09 | 0.16 | 0.35 | 0.57 | 0.24 | 2.67 |
| Cfer | 0.02 | 0.05 | 0.07 | 0.05 | 0.05 | 0.04 | 0.22 | 0.23 | 0.21 | 0.02 | 0.02 | 0.04 | 0.05 | 0.09 | 0.19 | 0.31 | 0.13 | 1.43 |
| Btau | 0.03 | 0.07 | 0.07 | 0.06 | 0.05 | 0.05 | 0.26 | 0.27 | 0.24 | 0.03 | 0.02 | 0.05 | 0.06 | 0.10 | 0.21 | 0.35 | 0.15 | 1.65 |
| Oorc | 0.03 | 0.09 | 0.11 | 0.09 | 0.08 | 0.07 | 0.38 | 0.40 | 0.35 | 0.04 | 0.03 | 0.07 | 0.09 | 0.14 | 0.31 | 0.52 | 0.22 | 2.41 |
| Fcat | 0.03 | 0.07 | 0.09 | 0.07 | 0.06 | 0.05 | 0.30 | 0.32 | 0.29 | 0.03 | 0.03 | 0.06 | 0.07 | 0.11 | 0.25 | 0.41 | 0.17 | 1.93 |
| Cfam | 0.03 | 0.08 | 0.09 | 0.07 | 0.07 | 0.06 | 0.32 | 0.34 | 0.31 | 0.03 | 0.03 | 0.06 | 0.07 | 0.12 | 0.27 | 0.44 | 0.19 | 2.05 |
| Mput | 0.01 | 0.01 | 0.02 | 0.01 | 0.01 | 0.01 | 0.07 | 0.07 | 0.06 | 0.01 | 0.01 | 0.01 | 0.01 | 0.03 | 0.05 | 0.09 | 0.04 | 0.42 |
| Hhya | 0.04 | 0.11 | 0.13 | 0.11 | 0.09 | 0.09 | 0.45 | 0.49 | 0.43 | 0.05 | 0.04 | 0.09 | 0.11 | 0.17 | 0.38 | 0.63 | 0.27 | 2.92 |
| Zcal | 0.01 | 0.03 | 0.03 | 0.03 | 0.02 | 0.02 | 0.10 | 0.11 | 0.09 | 0.01 | 0.01 | 0.02 | 0.02 | 0.04 | 0.08 | 0.13 | 0.06 | 0.63 |
| Eeur | 0.01 | 0.01 | 0.01 | 0.01 | 0.01 | 0.01 | 0.05 | 0.05 | 0.04 | 0.01 | 0.01 | 0.01 | 0.01 | 0.02 | 0.04 | 0.06 | 0.03 | 0.29 |
| Ccri | 0.02 | 0.05 | 0.07 | 0.05 | 0.05 | 0.04 | 0.21 | 0.23 | 0.20 | 0.02 | 0.02 | 0.04 | 0.05 | 0.08 | 0.17 | 0.29 | 0.13 | 1.37 |
| Sara | | 0.02 | 0.02 | 0.02 | 0.01 | 0.01 | 0.07 | 0.08 | 0.07 | 0.01 | 0.01 | 0.01 | 0.01 | 0.03 | 0.06 | 0.11 | 0.05 | 0.48 |
| Mmus | | | 0.06 | 0.05 | 0.04 | 0.04 | 0.21 | 0.22 | 0.19 | 0.02 | 0.02 | 0.04 | 0.05 | 0.08 | 0.17 | 0.28 | 0.12 | 1.31 |
| Moch | | | | 0.06 | 0.05 | 0.05 | 0.24 | 0.25 | 0.23 | 0.03 | 0.02 | 0.05 | 0.05 | 0.09 | 0.19 | 0.33 | 0.14 | 1.53 |
| Hgla | | | | | 0.04 | 0.03 | 0.19 | 0.21 | 0.18 | 0.02 | 0.02 | 0.04 | 0.04 | 0.07 | 0.16 | 0.27 | 0.11 | 1.23 |
| Ocun | | | | | | 0.03 | 0.17 | 0.17 | 0.15 | 0.02 | 0.02 | 0.03 | 0.03 | 0.06 | 0.13 | 0.23 | 0.09 | 1.05 |
| Opri | | | | | | | 0.15 | 0.16 | 0.14 | 0.01 | 0.01 | 0.03 | 0.03 | 0.05 | 0.13 | 0.21 | 0.09 | 0.96 |
| Hsap | | | | | | | | 0.85 | 0.76 | 0.08 | 0.07 | 0.15 | 0.17 | 0.31 | 0.67 | 1.11 | 0.47 | 5.15 |
| Ptro | | | | | | | | | 0.81 | 0.09 | 0.07 | 0.17 | 0.19 | 0.32 | 0.71 | 1.17 | 0.49 | 5.47 |
| Mfas | | | | | | | | | | 0.08 | 0.07 | 0.15 | 0.17 | 0.29 | 0.63 | 1.05 | 0.44 | 4.87 |
| Mmur | | | | | | | | | | | 0.01 | 0.01 | 0.02 | 0.03 | 0.07 | 0.11 | 0.05 | 0.53 |
| Ogar | | | | | | | | | | | | 0.01 | 0.01 | 0.03 | 0.06 | 0.10 | 0.04 | 0.46 |
| Dnov | | | | | | | | | | | | | 0.03 | 0.06 | 0.13 | 0.21 | 0.09 | 0.99 |
| Lafr | | | | | | | | | | | | | | 0.07 | 0.15 | 0.24 | 0.10 | 1.13 |
| Etel | | | | | | | | | | | | | | | 0.25 | 0.42 | 0.17 | 1.95 |
| Eedw | | | | | | | | | | | | | | | | 0.91 | 0.39 | 4.25 |
| Mdom | | | | | | | | | | | | | | | | | 0.64 | 7.09 |
| Oana | | | | | | | | | | | | | | | | | | 2.99 |
| Shar | | | | | | | | | | | | | | | | | | |

**Supplemental Fig. S8: The method to determine orthologous NUMTs between two species and the mathematical expectation of orthology assignments. a)** Schematic graphs showing the method of orthologous NUMT determination. NUMTs are regarded as orthologous between two species only if they are located in the same genomic synteny block within a distance of 6 protein-coding genes (3 genes upstream and 3 genes downstream), and their sequences overlap each other at least 50%. **b)** The simplified formula estimating the probability of two NUMTs that are assigned as orthologs by chance. Suppose that the average number of protein-coding genes in mammalian genomes is 20,000, that the average length of NUMTs is 200 bp, and that the average size of mammalian mitogenomes is 16,600 bp. The error rate was calculated at $3.03 \times 10^{-6}$. **c)** The mathematic expectations of orthology assignment error. For each pair of species, the mathematic expectation was calculated by multiplying the error rate ($3.03 \times 10^{-6}$) by all possible NUMT pairs between these two species. The heatmap indicates that the error expectation is smaller than 1 for most comparisons except the comparisons between *S. harrisii* and other species.

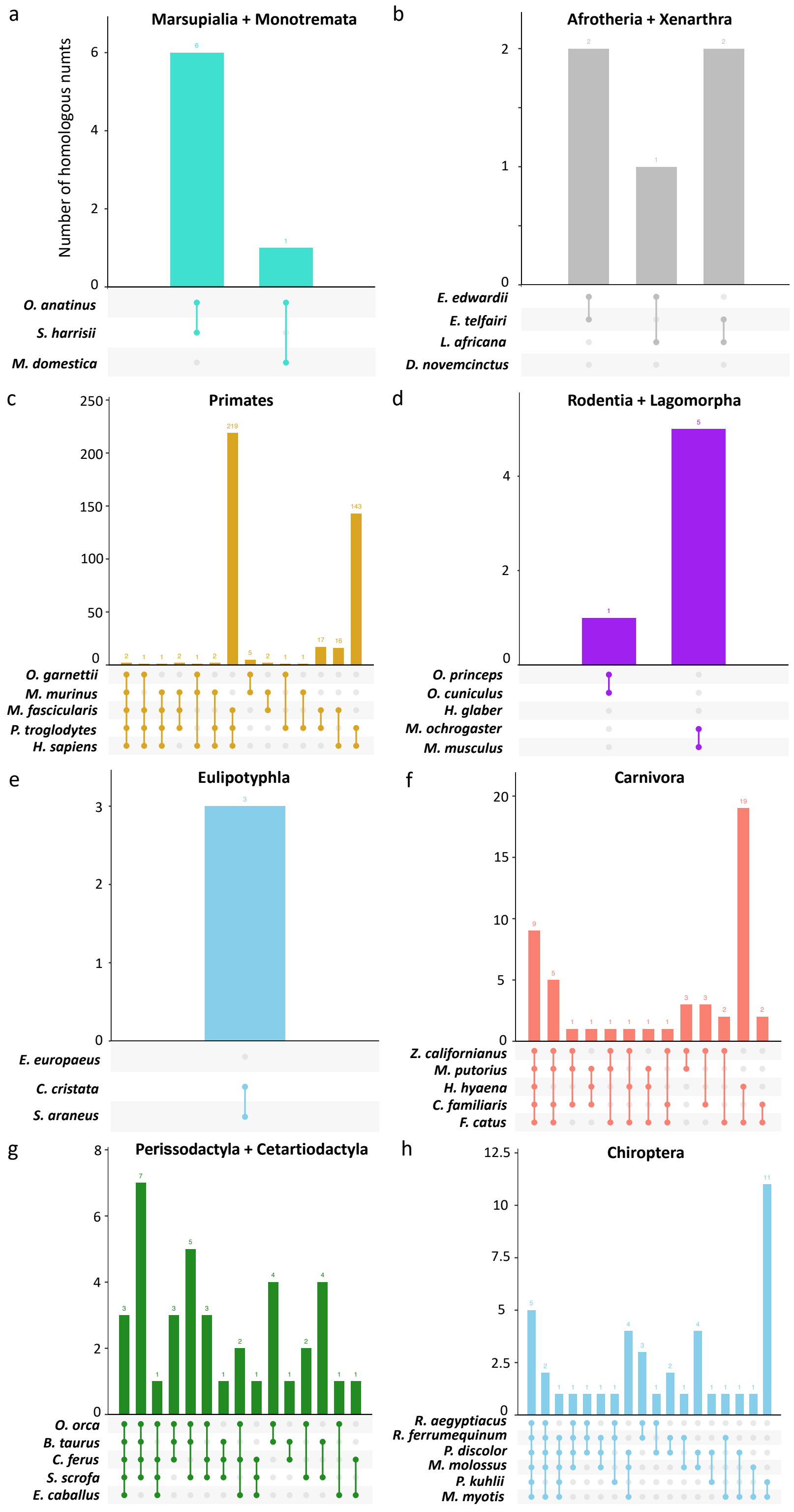


**Supplemental Fig. S9: The UpSetR plots showing the number of homologous NUMTs that were predicted to be shared amongst species in the eight defined clades.** In each panel, the bars above the vertical lines connecting different species indicate the number of homologous NUMTs detected amongst these species. **a)** Marsupialia and Monotremata (n = 3); **b)** Afrotheria and Xenarthra (n = 4); **c)** Primates (n= 5); **d)** Rodentia and Lagomorpha (n = 5); **e)** Eulipotyphla (n = 3); **f)** Carnivora (n = 5); **g)** Perrisodactyla and Cetartiodactyla (n = 5); **h)** Chiroptera (n = 6).

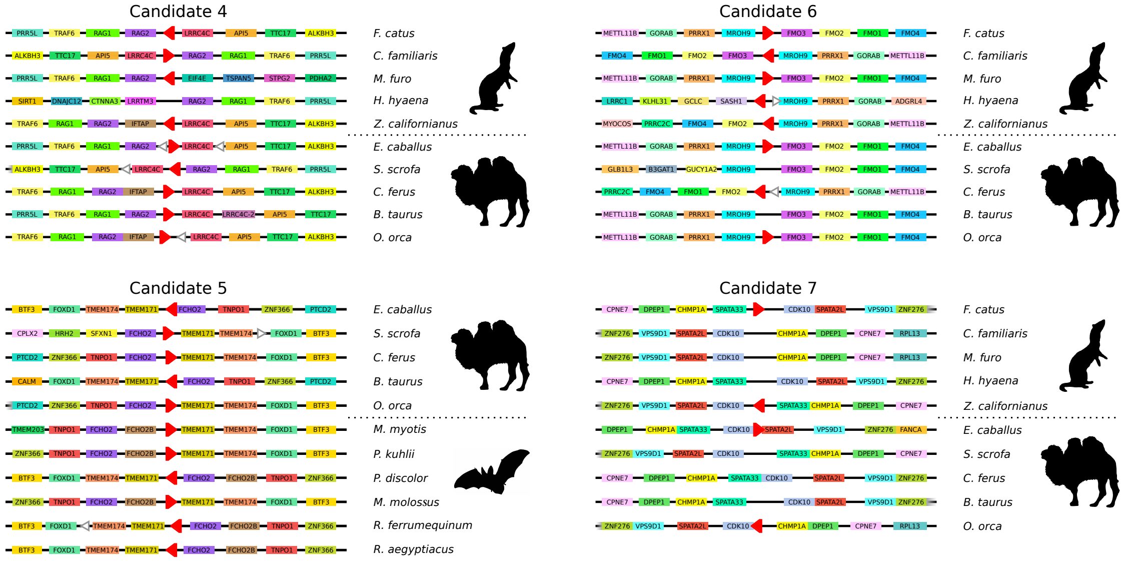


**Supplemental Fig. S10: Visualisation of the genomic microsynteny of 4 ancestral NUMT blocks across the defined clades.** For each candidate, the lines represent the conserved microsynteny blocks across species. Protein-coding genes are shown as rectangles on the lines, which are colour-coded. Genes with no gaps between each other indicate that they have overlapping genomic coordinates. NUMTs are represented by triangles, with its direction relative to the corresponding mtDNA indicated. Red triangles represent ancestral orthologous NUMTs, while white ones represent NUMTs that are considered as non-orthologous. The triangles that overlap with protein-coding genes indicate that the NUMTs are located in the intronic regions of these protein-coding genes; otherwise, the NUMTs are located in intergenic regions or 3'-UTRs.

**Supplemental Table S8:** The combinations of Megablast parameters used to optimise the NUMT identification pipeline.

| Combinations of Parameters | Number of numts predicted in hg38 | average identity (%) | average length (bp) |
|---|---|---|---|
| default blastn | 186 | 85.35 | 1431.01 |
| -template_type optimal -template_length 18 -word_size 11 | 846 | 80.87 | 641.78 |
| -template_type optimal -template_length 16 -word_size 11 | 845 | 80.88 | 641.06 |
| -template_type optimal -template_length 21 -word_size 11 | 845 | 80.87 | 642.50 |
| -template_type coding -template_length 18 -word_size 11 | 841 | 80.84 | 645.18 |
| -template_type coding_optimal -template_length 18 -word_size 11 | 844 | 80.84 | 643.29 |
| -template_type optimal -template_length 18 -word_size 12 | 839 | 80.92 | 645.78 |
| -template_type optimal -template_length 16 -word_size 12 | 835 | 80.93 | 648.26 |
| -template_type optimal -template_length 21 -word_size 12 | 836 | 80.89 | 648.26 |
| -template_type coding -template_length 18 -word_size 12 | 819 | 80.93 | 658.37 |
| -template_type coding_and_optimal -template_length 18 -word_size 12 | 841 | 80.86 | 644.96 |

**Supplemental Table S9:** The optimal substitution models according to Bayesian Information Criterion (BIC) in ModelFinder used in IQTREE.

| Trees | Length (bp) | Model | BIC |
|---|---|---|---|
| CYTB | 1472 | GTR+F+G4 | 176928.46 |
| ND1 | 1174 | K3Pu+F+R6 | 159998.47 |
| Candidate 1 | 228 | HKY+F+I | 2403.98 |
| Candidate 2 | 342 | TPM3+F | 3258.08 |
| Candidate 3 | 253 | TPM2u+F | 1859.05 |

**Supplemental Table S10:** The RNA-Seq samples used for NUMT expression analyses.

| Taxon | Species | Tissue | SRA Accession |
|---|---|---|---|
| Monotremata | *Ornithorhyncus anatinus* | brain | SRR5412225 |
| Monotremata | *Ornithorhyncus anatinus* | brain | SRR5412222 |
| Monotremata | *Ornithorhyncus anatinus* | liver | SRR5412236 |
| Monotremata | *Ornithorhyncus anatinus* | liver | SRR5412234 |
| Monotremata | *Ornithorhyncus anatinus* | kidney | SRR5412233 |
| Monotremata | *Ornithorhyncus anatinus* | kidney | SSR5412232 |
| Monotremata | *Ornithorhyncus anatinus* | heart | SSR5412227 |
| Monotremata | *Ornithorhyncus anatinus* | heart | SSR5412229 |
| Rodentia | *Heterocephalus glaber* | brain | SSR1959171 |
| Rodentia | *Heterocephalus glaber* | brain | SSR1959162 |
| Rodentia | *Heterocephalus glaber* | kidney | SSR5517229 |
| Rodentia | *Heterocephalus glaber* | kidney | SSR5517227 |
| Rodentia | *Heterocephalus glaber* | liver | SSR5517452 |
| Rodentia | *Heterocephalus glaber* | liver | SSR5517450 |
| Rodentia | *Heterocephalus glaber* | heart | SSR5517306 |
| Rodentia | *Heterocephalus glaber* | heart | SSR5517305 |
| Primates | *Home Sapiens* | brain | SRR8942877 |
| Primates | *Home Sapiens* | brain | SRR8942874 |
| Primates | *Home Sapiens* | kidney | SRR12937853 |
| Primates | *Home Sapiens* | kidney | SRR12937852 |
| Primates | *Home Sapiens* | liver | SRR15694280 |
| Primates | *Home Sapiens* | liver | SRR15694281 |
| Primates | *Home Sapiens* | heart | SRR19552233 |
| Primates | *Home Sapiens* | heart | SRR19552232 |
| Carnivora | *Canis familiaris* | brain | SSR12899123 |
| Carnivora | *Canis familiaris* | brain | SSR12899118 |
| Carnivora | *Canis familiaris* | kidney | SSR19614474 |
| Carnivora | *Canis familiaris* | kidney | SSR19614479 |
| Carnivora | *Canis familiaris* | liver | SSR19614475 |
| Carnivora | *Canis familiaris* | liver | SSR19614488 |
| Carnivora | *Canis familiaris* | heart | SSR8474294 |
| Carnivora | *Canis familiaris* | heart | SSR8474261 |
| Chiroptera | *Rousettus aegyptiacus* | brain | SRR2913353 |
| Chiroptera | *Rousettus aegyptiacus* | brain | SRR2914295 |
| Chiroptera | *Rousettus aegyptiacus* | kidney | SRR2914360 |
| Chiroptera | *Rousettus aegyptiacus* | kidney | SRR2913355 |
| Chiroptera | *Rousettus aegyptiacus* | liver | SRR2914369 |
| Chiroptera | *Rousettus aegyptiacus* | liver | SRR2914059 |
| Chiroptera | *Rousettus aegyptiacus* | heart | SRR2914359 |
| Chiroptera | *Rousettus aegyptiacus* | heart | SRR2913354 |
| Cetartiodactyla | *Bos taurus* | brain | SRR11657443 |
| Cetartiodactyla | *Bos taurus* | brain | SRR11657439 |
| Cetartiodactyla | *Bos taurus* | kidney | SRR19745555 |
| Cetartiodactyla | *Bos taurus* | kidney | SRR19745554 |
| Cetartiodactyla | *Bos taurus* | liver | SRR15721701 |
| Cetartiodactyla | *Bos taurus* | liver | SRR15721700 |
| Cetartiodactyla | *Bos taurus* | heart | SRR12442762 |
| Cetartiodactyla | *Bos taurus* | heart | SRR12442763 |
